# CLASP1 is essential for neonatal lung function and survival in mice

**DOI:** 10.1101/2022.04.27.489792

**Authors:** Ana L. Pereira, Tiago F. da Silva, Luísa T. Ferreira, Martine Jaegle, Marjon Buscop-van Kempen, Robbert Rottier, Wilfred F. J. van Ijcken, Pedro Brites, Niels Galjart, Helder Maiato

## Abstract

The first breath of air at birth marks the beginning of extrauterine life, and breathing problems due to incomplete lung development or acute respiratory distress are common in premature babies and respiratory diseases. However, the underlying molecular mechanisms remain poorly understood. Here we show that the microtubule plus-end-tracking protein CLASP1 is required for neonatal lung function and survival. CLASP1 is expressed in the lungs and associated respiratory structures throughout embryonic development. *Clasp1* disruption in mice caused intrauterine growth restriction and neonatal lethality due to acute respiratory failure. Knockout animals showed impaired lung inflation associated with smaller rib cage formation and abnormal diaphragm innervation. Live-cell analysis of microtubule dynamics in cultured hippocampal neurons revealed an increased catastrophe rate, consistent with a role of CLASP1 in neurite outgrowth. Histological and gene expression studies indicated that CLASP1 is required for normal pneumocyte differentiation and fetal lung maturation. Thus, CLASP1-mediated regulation of microtubule dynamics assists multiple systems essential for neonatal lung function and survival.

## Introduction

CLIP-Associating Proteins (CLASPs) are microtubule (MT) plus-end-tracking proteins with important cellular roles in the regulation of MT dynamics (Lawrence et al., 2020). In yeast and *Drosophila*, the single *Clasp* orthologue is essential for mitosis and viability (Grallert et al., 2006; Inoue et al., 2000; Lemos et al., 2000; Pasqualone and Huffaker, 1994b). *C. elegans* have three *Clasp* orthologues, but only one is essential for viability (Cheeseman et al., 2005). Mammals have two paralogues, *Clasp1* and *Clasp2*, which play partially redundant roles during cell division and migration *in vitro* (Akhmanova et al., 2001; Drabek et al., 2006; Inoue et al., 2000; Lemos et al., 2000; Maiato et al., 2003a; Mimori-Kiyosue et al., 2005; Mimori-Kiyosue et al., 2006; Pereira et al., 2006). Mammalian CLASP2 is predominantly expressed in the brain, reproductive system and blood cells (Akhmanova et al., 2001; Nagase et al., 2000). In agreement, *Clasp2* knockout (KO) mice are sterile and, if they do not succumb to hemorrhages associated with compromised hematopoiesis, they can live well into adulthood (Drabek et al., 2012). Mammalian CLASP1 is more ubiquitously expressed (Akhmanova et al., 2001; Nagase et al., 2000), suggesting a broader, possibly more essential, physiological role, which has so far not been determined.

In addition to their canonical localization at growing MT plus-ends (Akhmanova et al., 2001), mammalian CLASPs are components of fundamental mitotic structures, including kinetochores (KTs), centrosomes, spindle midzone and midbody, and play important roles in chromosome congression and segregation, maintenance of spindle bipolarity and regulation of MT dynamics at KTs (Girao et al., 2020; Inoue et al., 2000; Lemos et al., 2000; Logarinho et al., 2012; Maffini et al., 2009; Maiato et al., 2003a; Maiato et al., 2005; Maiato et al., 2003b; Maiato et al., 2002; Pereira et al., 2006). In interphase, CLASPs also associate with the trans-Golgi network via molecular and functional interactions with GCC185 (Efimov et al., 2007), and participate in generating polarized microtubule networks (Akhmanova et al., 2001; Drabek et al., 2006; Wittmann and Waterman-Storer, 2005), as well as in the stabilization and rescue of microtubule plus-ends at their cortical attachment sites (Mimori-Kiyosue et al., 2005), instrumental for processes such as wound healing-induced fibroblast motility (Drabek et al., 2006). Moreover, CLASPs are required for growth cone orientation and axon guidance, suggesting a specialized role in neuronal cell biology (Hur et al., 2011; Lee et al., 2004; Sayas et al., 2019). In agreement, CLASP2 has been implicated in neuron migration, morphogenesis, cortical lamination (Dillon et al., 2017), as well as in microtubule capture at neuromuscular junctions (Schmidt et al., 2012).

Mammalian lungs consist of a highly branched network containing thousands to millions of airways arrayed in intricate patterns essential for respiration. In mice, the lungs arise from the foregut endoderm at embryonic day 9.5 (E9.5) (Whitsett et al., 2019). At this stage, lung buds undergo repeated rounds of branching and outgrowth. During the pseudoglandular stage (∼E10.5-E14.5), the conductive airways are formed and lined with a mixture of secretory (club cells), neuroendocrine (NE) and basal cells (Whitsett et al., 2019). The distal buds of the lungs then elongate at the cannalicular stage (∼E14.5-E16.5) and ultimately give rise to the terminal sacs containing type I and II epithelial cells (aka pneumocytes) (Whitsett et al., 2019; Whitsett et al., 2010). The saccular stage, which occurs between E17.5 to the fifth day of postnatal development in mice, is a particular lung morphogenesis stage in which the lungs switch from fluid-filled to air-filled structures, upon which postnatal survival depends. This stage is marked by dilatation of peripheral airspaces, differentiation of the respiratory epithelium, increased vascularity and surfactant synthesis. Undifferentiated cuboidal cells turn into type I and II pneumocytes. Type II pneumocytes contain lamellar bodies, which are intracellular storage units of surfactant, whose secretion is essential for normal lung development and function after birth (Guo et al., 2019). Type II pneumocytes are thought to give rise to type I pneumocytes (Barkauskas et al., 2013; Desai et al., 2014; Nabhan et al., 2018), which line the greater part of the alveolar surface and are responsible for gas-exchange. Postnatally, the primitive saccules enlarge and are subdivided into smaller units by a septation process called “alveolization” (Maeda et al., 2007; Warburton et al., 2006). Alveolization greatly increases the surface area and lung compliance, and occurs in late gestation in humans and postnatally in mice. The pulmonary surfactant system is one of the latest systems developed during lung maturation and is composed by phospholipids (80%), neutral lipids (12%), and proteins (8%). The protein fraction of the surfactant system is made by surfactant protein – A (SP-A), B (SP-B), C (SP-C) and D (SP-D). The most important surfactant proteins for life after birth are SP-B and SP-C. The surfactant is secreted to the alveolar lumen and coats the intra-alveolar wall, decreasing the tension at the air-liquid interface that facilitates the expansion of the alveoli during respiration (Boyden, 1977).

The biggest challenge of extrauterine life is the first breath of air after lung activation at birth. In mice, when pups are born, they become cyanotic in the first couple of minutes, between ceasing of the placental oxygen support by the mother and the time that breathing takes over after lung function activation, from which normal pups recover their pinkish color. However, any abnormality that affects different physiological systems that support breathing can be life-threatening. Partial or total lung collapse is one of the most common hallmarks of respiratory diseases and it may occur in the absence of a primary lung problem, due to cardiovascular, neuromuscular and/or skeletal deficiencies, among others (Turgeon and Meloche, 2009). Lung collapse is usually related to insufficient maturation of the lung tissue after delivery. In humans, the most common cause of pulmonary syndrome distress in newborns relates with insufficient synthesis of surfactant or genetic mutations in surfactant proteins (Nkadi et al., 2009). In these cases, type II pneumocytes may not be sufficiently matured, and surfactant synthesis and secretion may be compromised to allow proper lung expansion. Some genes implicated in lung maturation lead to lung collapse upon disruption in mice. For instance, *Nfib, Pdpn* and *Ndst1* knockout mice, despite visible efforts to breathe, show insufficient inflation for proper gas-exchange and die within few minutes after birth. *Pdpn* deficient mice have abnormal distal saccules with blockade in type I pneumocyte differentiation. However, surfactant levels are high, which indicates that maturation of type II pneumocytes occurs normally (Ramirez et al., 2003). In contrast, type II pneumocyte differentiation seems to be compromised in *Ndst1* deficient animals, resulting in impaired surfactant production (Fan et al., 2000).

Here we investigate the physiological roles of CLASP1 after targeted disruption of the *Clasp1* gene in mice. Although *Clasp1* disruption led to intrauterine growth restriction, CLASP1 was mostly dispensable for embryonic development in mice, consistent with previously uncovered redundancy with CLASP2 during cell division. Surprisingly, we found that CLASP1 is required immediately after birth for neonatal lung function and survival. Histological and functional analyses in *Clasp1* knockout (KO) animals and cultured hippocampal neurons, complemented by gene expression studies in the lungs, unveiled an unexpected role of CLASP1-mediated control of microtubule dynamics in the integration of growth control, lung cell differentiation and normal respiratory muscle innervation, essential to support neonatal lung function and survival after birth.

## Results

### Clasp1 disruption leads to intrauterine growth restriction and neonatal lethality in mice

To gain insight into the physiological role of CLASP1 in mammals, we started by investigating its expression *in situ* by immunohistochemistry (IHC) in late embryonic tissues. We found that CLASP1 is abundantly expressed in the endothelium of blood vessels, as well as in secretory cells of mouse lungs at E18.5 (Fig. 1A). CLASP1 is also expressed in other major respiratory structures, such as the diaphragm (the primary respiratory muscle) and intercostal muscles associated with the rib cage, as well as in other organs, including the heart, the adrenal gland, and the kidneys (Fig. 1A).

**Figure 1.**
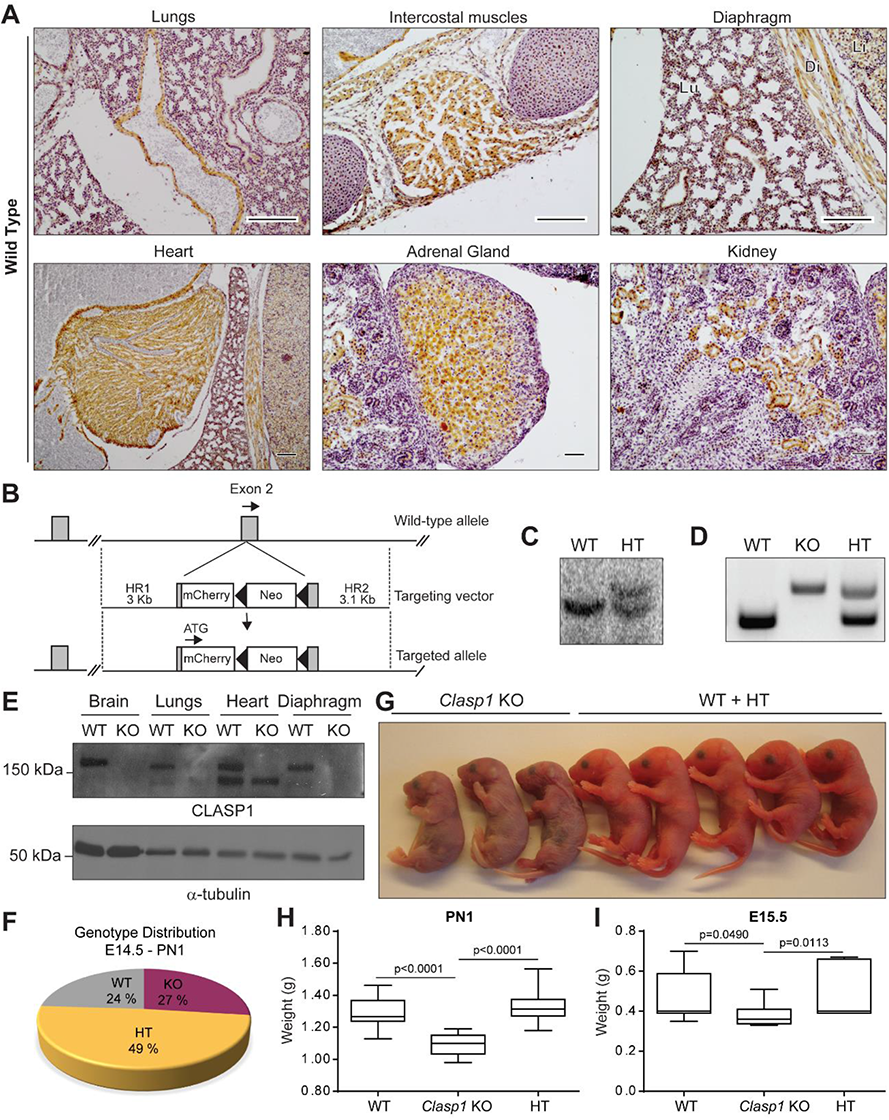
Targeted inactivation of the *Clasp1* allele. (A) Multi-organ systematic analysis of CLASP1 expression by immunohistochemistry in WT mice. CLASP1 was expressed in the endothelium, secretory and ciliated cells in lungs of WT embryos. The protein can also be detected in associated respiratory tissues, including the diaphragm and intercostal muscles, as well as in the heart, adrenal glands and kidneys. (B) Schematic representation of the targeting strategy. The second *Clasp1* exon was interrupted by insertion of a mCherry-loxP-pMC1NEO-loxP cassette. The neomycin resistance gene (neo) is transcribed in the opposite direction of *Clasp1*. LoxP sites are indicated by arrowheads. The targeted allele is represented at the bottom. (C) Southern blot analysis of targeted embryonic stem (ES) cell DNA – a positive clone for homologous recombination shows both wild-type and KO alleles. (D) PCR of genomic DNA extracted from wild-type and *Clasp1* KO tails of newborns (postnatal day 1). The lower band corresponds to the wild-type allele, while the higher band is consistent with the targeted KO allele. (E) Western blot analysis of CLASP1 in embryonic organs collected at E18.5 from WT and KO embryos. α-tubulin was used as loading control. (F) Global analysis of genotype distribution of Clasp1 WT (n=111), HT (n=225) and KO (n=126) animals from embryonic day 14.5 until postnatal day 1 (E14.5-PN1) shows that mutant mice develop and are born at the expected Mendelian ratio. (G) Representative photographic pictures of cyanotic *Clasp1^-/-^* newborns and WT littermates. (H, I) Body weight distribution of the WT and KO animals is shown at PN1 (H) and E15.5 (I). Groups consisted of (H) 15 WT, 21 HT and 19 KO embryos, and (I) 8 WT, 10 KO and 7 HT.

To investigate the physiological role of CLASP1 in mammals we disrupted the murine *Clasp1* gene by insertion of a mCherry-loxP-pMC1neo-loxP cassette in the second exon, adjacent to the beginning of the *Clasp1* coding region, using homologous recombination in embryonic stem cells. In this modified *Clasp1* allele, the expression of CLASP1 was interrupted by the transcription of the mCherry open reading frame and by the transcription of the neomycin resistance gene (neo), which is antisense with respect to the *Clasp1* gene (Fig. 1B). Chimeric mice transmitted the mutant *Clasp1* allele to the germ line, as determined by Southern blot and PCR analyses (Fig. 1C, D). The ablation of CLASP1 expression was confirmed by western blot analysis using multiple organs collected at E18.5 from WT and KO embryos (Fig. 1E). Interestingly, in the heart, a smaller CLASP1 isoform is still expressed in both WT and KO animals, despite the insertion cassette in exon 2.

In an effort to generate *Clasp1* KO mice, *Clasp1* HT females and males were repeatedly intercrossed. However, genotyping of litters derived from the intercrossing failed to identify any homozygous mutant (KO) offspring, with only WT and HT mice (both males and females) being obtained. Careful analysis and monitoring of timed pregnancies demonstrated that *Clasp1*-null embryos were viable until the end of gestation and birth, and were born at near-Mendelian ratios (Fig. 1F). Thus, homozygous *Clasp1* disruption did not impair *in utero* survival in mice. Nevertheless, after birth, all *Clasp1^-/-^* pups stopped responding to pinching stimuli within 20-30 min after delivery and died following various concerted gasping movements in a visible effort to breathe. Contrary to WT and HT pups that initiated normal respiration, acquired a pink color, and rapidly inflated their lungs shortly after birth, all postnatal KO pups (n=20) died presenting cyanotic features, an indication of severe respiratory distress (Fig. 1G). Additionally, *Clasp1*^-/-^ pups exhibited an evident reduction in fetal growth (Fig. 1G). At postnatal day 1 (PN1), total body weight of the KO pups was reduced almost 20% relative to WT or HT littermates: *Clasp1^+/+^* 1.291 ± 0.097g *vs. Clasp1^+/-^* 1.328 ± 0.087g *vs. Clasp1^-/-^* 1.086 ± 0.069g, p≤0.0001 relative to both WT and HT, Mann-Whitney test (Fig. 1H). This reduction in body weight was already visible at E15.5 (Fig. 1I). Thus, homozygous targeting of *Clasp1* prevents normal fetal growth and causes postnatal lethality, likely due to respiratory failure.

### Loss of CLASP1 results in a smaller ribcage and a slight delay in ossification

To investigate whether the observed respiratory distress in *Clasp1* KO mice were a direct consequence of skeleton abnormalities associated with their reduced body weight we performed whole-mount cartilage and mineralized bone staining with alcian blue and alizarin red. Despite being smaller overall, including evident microcephaly with smaller orbits, late embryonic skeletons derived from *Clasp1* KO mice (E18.5) were complete and without any gross abnormalities (Fig. 2A,B). However, careful inspection revealed a smaller rib cage, with slightly delayed ossification in specific areas, such as in the sternum and occipital bones (Fig. 2C-C’’), consistent with immature development and intra-uterine growth restriction (IUGR).

**Figure 2.**
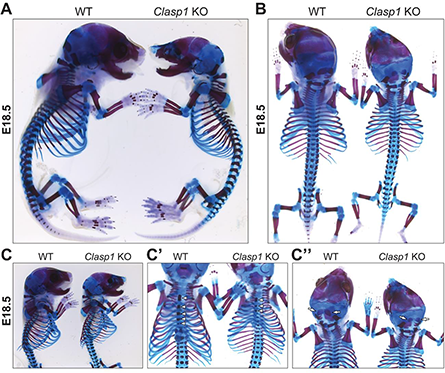
*Clasp1* KO mice show microcephaly, reduced rib-cage and delayed ossification. Alcian blue and alizarin red staining of E18.5 littermates. Lateral (A) and dorsal (B) view of the animals show clear differences in size and volume of the head and thoracic cavity in *Clasp1* KO mice compared to WT. Lateral (C), ventral (C’) and dorsal (C’’) views of the rib cage. The mutant shows a small delay in sternum ossification (arrowheads) and in the occipital bone (arrows).

### Clasp1^-/-^ mouse embryonic fibroblasts show incidental cell division defects

To investigate possible underlying causes of IUGR in *Clasp1* KO animals, we performed immunofluorescence (IF) and western blot analyses on MEFs derived from WT and KO embryos collected at day E11.5 (Fig. S1A, B). This analysis revealed that E11.5 *Clasp1* KO MEFs displayed an abnormally high percentage of mitotic cells with multipolar spindles (∼20%) and supernumerary centrosomes (∼31%) (Fig. S1A, C). The presence of bi-nucleated cells (possibly as a consequence of cytokinesis failure) in the *Clasp1* KO MEFs was also noticed (Fig. S1D), which explains the observed increase in mitotic cells with multipolar spindles and supernumerary centrosomes. Curiously, CLASP2-deficient keratinocytes have also been reported to exhibit increased centrosomal numbers and multipolar spindles that prevent cell proliferation in a p53-dependent manner (Shahbazi et al., 2017). Since multipolar mitosis are normally associated with tetraploidy induced by cytokinesis failure, which is known to cause a p53-dependent cell cycle arrest (Kuffer et al., 2013), the cell division defects observed in MEFs help to explain the smaller size of *Clasp1* KO animals associated with IUGR.

### Clasp1 KO mice have an abnormal innervation pattern of the diaphragm

Respiratory failure may also be caused by anomalies in the nervous system that feed into associated muscles. Therefore, we next investigated the innervation pattern of the phrenic nerve in diaphragms from WT and *Clasp1* KO embryos. In order to visualize innervation in whole-mounts of diaphragms from E14.5-E18.5, we immunolabeled muscles with antibodies against βIII-tubulin and synaptic vesicle glycoprotein 2A (SV2). We found that *Clasp1* KO mice showed an abnormal pattern characterized by decreased branching of the phrenic nerve from E14.5 to E18.5 but with larger extensions that covered a wider innervated area within the muscle (Fig. 3A-C). To determine whether the abnormal innervation observed in *Clasp1* KO mice affected the formation and localization of the neuromuscular junctions (NMJs), we labelled these synapses with α-bungarotoxin. *Clasp1* KO mice displayed an increased dispersion of NMJs (Fig. 3D), consistent with the increased nerve extension (Fig. 3C). However, *Clasp1* KO mice showed a synaptic defect, with reduced number and density of NMJs (Fig. 3D,E). Importantly, the sarcomere ultra-structure and diaphragm morphology was unaltered in *Clasp1* KO mice (Fig. S2). Combined, these results indicate that the loss of *Clasp1* impacts the ability of motor neurons to innervate the diaphragm.

**Figure 3.**
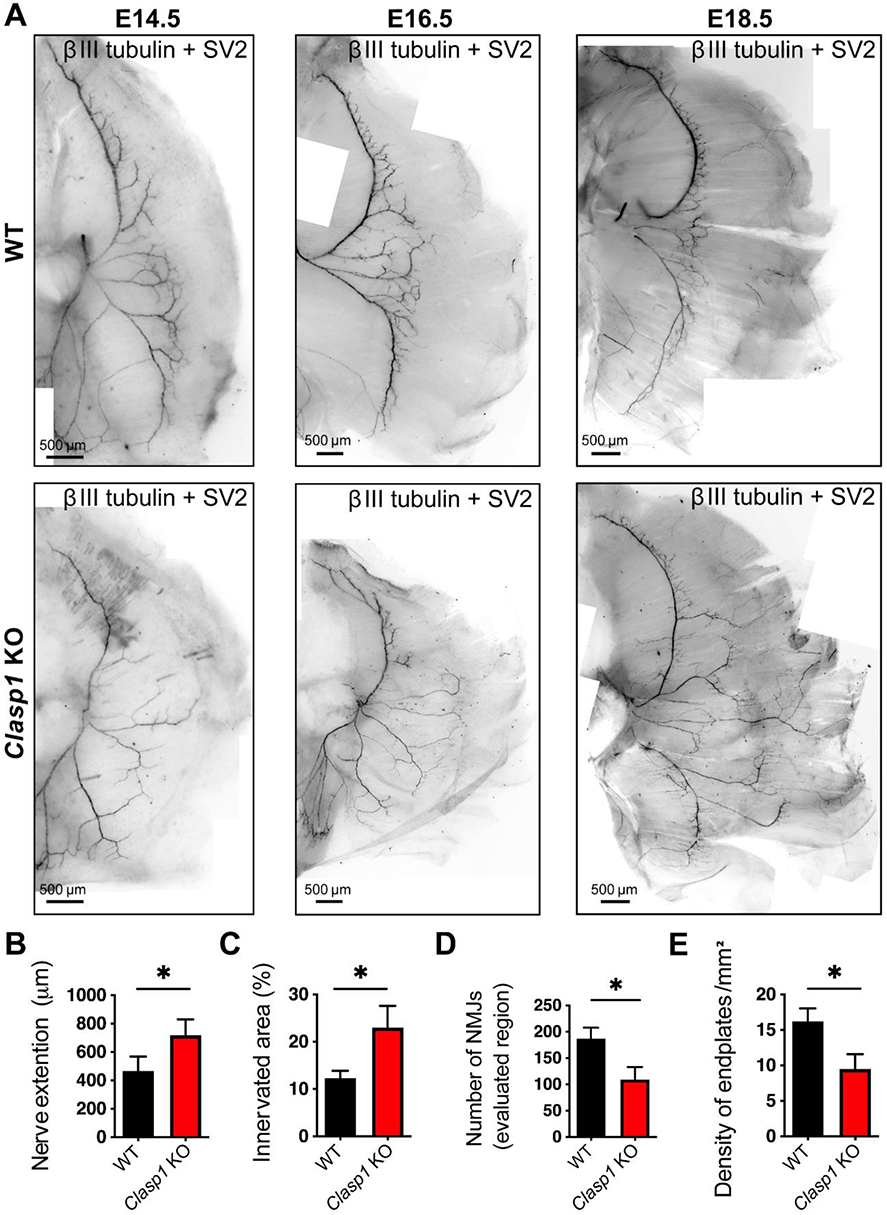
Innervation of the diaphragm is impaired in *Clasp1* KO mice from E14.5-E18.5. (A) A cocktail of antibodies composed of β-III tubulin and SV2 was used to label the motor nerve and pre-synaptic terminals in the diaphragm at different developmental stages. Scale bar 500 μm. (B) The measurement of the motor projections extending from the main nerve trunk reveal an increased travelled distance in the *Clasp1* KO when compared to the WT. The labelling of the diaphragms at E18 with BGTx revealed an increased innervated region (C) and a decreased number of synapses (D) in the *Clasp1* KO which lead to a decreased density of endplates (E).

### Clasp1 KO neurons show altered growth capacity in vitro

The abnormalities in the patterned innervation of the diaphragm prompted us to investigate whether the loss of CLASP1 affects the growth of axons. Motor neurons (MNs) were isolated from WT and *Clasp1* KO E12.5 embryos and plated onto matrigel, or PLO+laminin coated glass slides. In these two conditions, WT MNs exhibited robust growth of axons and dendrites at 16 h (Fig. 4A,B) and 48 h (Fig. 4A,C) after plating. However, MNs from *Clasp1* KO mice displayed decreased axon and dendrite growth (Fig. 4A-C). To determine whether the defect in axon growth was intrinsic to the loss of CLASP1 function, we determined whether other neuron types also displayed defects in neurite growth. Dorsal root ganglion (DRGs) neurons from E12.5 embryos of WT and *Clasp1* KO mice were isolated, cultured and neurite growth was evaluated. Similar to MNs, DRG neurons from *Clasp1* KO mice displayed a defect in neurite growth (Fig. 4D). The length of the longest neurite (Fig. 4E), as well as the total length of all neurites (Fig. 4F) was smaller in *Clasp1* KO DRG neurons when compared to WT DRG neurons. Together, these results highlight the importance of CLASP1 for normal axon growth.

**Figure 4.**
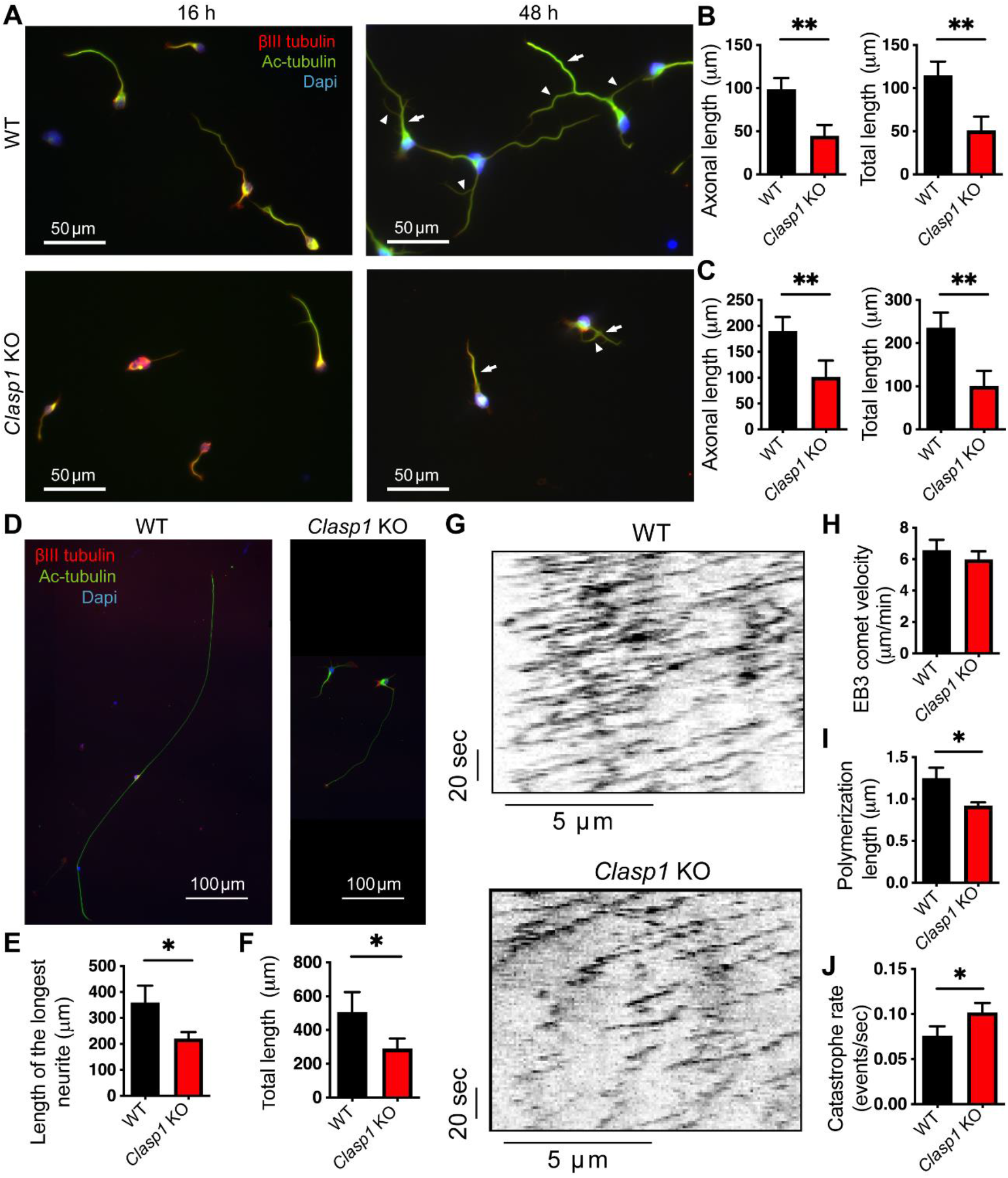
Neurite outgrowth, branching capacity and microtubule dynamics are altered in *Clasp1* KO neurons. (A) Fixed and immunostained MNs (E12.5) with acetylated tubulin (red), β-III tubulin (green) and DAPI (blue). Scale bar 50 μm. Neurite outgrowth evaluation show a decreased capability of *Clasp1* KO MNs to extend the axon (arrows) and dendrites (arrowheads) to a length similar to the one found in WT MNs at 16 h (B), and 48 h (C). (D) Fixed and immunostained DRG neurons with acetylated tubulin (red), β-III tubulin (green) and DAPI (blue). Scale bar 100 μm. Similar to MNs, *Clasp1* KO DRG neurons show a decrease capability for the extension of the longest neurite (E) and total neurite length (F). (G) Kymograph analysis of EB3-GFP comets in WT and *Clasp1* KO hippocampal neurons (E18.5) in culture. The velocity of EB3-GFP comets is similar in both WT and KO neurons (H). Polymerization length is decreased in *Clasp1* KO neurons (I), which show an increased catastrophe rate relative to WT (J).

In order to identify possible alterations in microtubule dynamics caused by the loss of CLASP1, we performed live-cell imaging of microtubule growth in cultured hippocampal neurons. Neurons from WT and *Clasp1* KO mice were transfected with a plasmid expressing EB3-GFP, a bonafide +TIP and fluorescent marker of microtubule growth behaviour (Stepanova et al., 2003). Analyses of EB3-GFP comets demonstrated that their velocity was similar in WT and *Clasp1* KO neurons (Fig. 4G,H). However, the polymerization length was significantly reduced in *Clasp1* KO neurons, whereas catastrophe rate was increased (Fig. 4I,J), consistent with a role for CLASP1 as a microtubule growth promoting factor. These results emphasize the consequences of CLASP1 loss in deregulation of microtubule dynamics and underscore the *in vivo* relevance of CLASP1 towards normal axon extension and establishment of synapses.

### CLASP1 is required for lung maturation and function in newborn mice

To directly examine lung function in the *Clasp1* KO mice, we collected the lungs from WT and KO newborn animals and analyzed their gross morphology. Lungs in KO mice lacked the typical foamy aspect of WT lungs (Fig. S3). To test for the presence of air, the lungs were placed in a test tube with phosphate-buffered saline solution (PBS) to monitor their floatation capacity. While the WT lungs floated on top of the saline solution, the KO lungs immediately sank to the bottom of the tube (Fig. S3), consistent with the absence of air in the lungs and the incapacity of the KO pups to breathe.

Despite the respiratory distress observed in KO neonates, *Clasp1* disruption did not interfere with the presence of expected cellular components of the mammalian lungs, including smooth muscle actin (SMA), cilia and club cells, as identified with antibodies against SMA, acetylated tubulin and α-CC10, respectively (Fig. S4A-C). We therefore sought to obtain a thorough perspective of lung morphology in *Clasp1* KO animals throughout embryonic development. To do so, we collected lungs from both WT and KO embryos and neonate littermates to perform comparative histological and morphological analyses. In general, despite being smaller, organogenesis and overall morphology seemed to be preserved in lungs from newborn *Clasp1* KO animals, with four right lobes and a single left lobe adjoining the heart, indicating that patterning during early lung development was not affected (our unpublished observations). Nevertheless, evaluation of hematoxylin and eosin (HE) sequential lung sections from KO neonatal pups revealed extensive and severe atelectasis (Fig. 5A) as the most likely cause of death, consistent with our observations of respiratory distress and cyanosis in *Clasp1* KO newborns.

**Figure 5.**
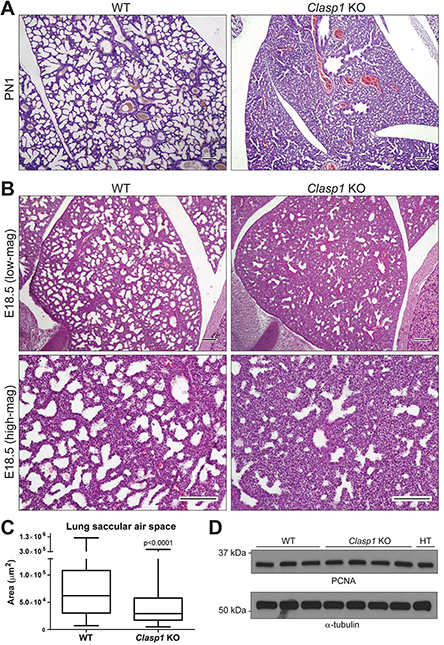
Histological and immunological examination of the *Clasp1* KO lungs demonstrating decreased air space. (A) Lung sections of postnatal day 1 in WT and *Clasp1*-null mice stained with HE demonstrate an extensive lung collapse in the *Clasp1* KO animals after failed attempts to breathe. (B) Histological sections in lower (upper panels) and higher (lower panels) magnifications of E18.5 lungs of WT and *Clasp1*-null mice stained with HE. Scale bars 150 µm. (C) Morphometric quantification of lung saccular airspace in WT and KO animals at E18.5. n=3 for all genotypes. A minimum of 3 different areas per slide were quantified. (D) Western blot analysis showing similar expression of PCNA as a proxy for cell proliferation in all the genotypes. α-tubulin was used as loading control.

To distinguish between atelectasis and hypercellularity, we studied fixed lungs from WT and KO E18.5 littermates, which were sacrificed before having had the chance to breathe. Both qualitative observation and quantitative morphometric measurements in three different animals revealed diminished saccular air space in the KO lungs (Fig. 5B, C), with a consequent increase in the amount of mesenchymal tissue, when compared to WT. Hypercellularity due to more active cell proliferation was excluded, as we found no difference in proliferating cell nuclear antigen (PCNA) levels between WT and KO lungs after western blot analysis (Fig. 5D). Moreover, immunostaining of lung sections with an antibody against cleaved caspase 3 revealed only a residual and equivalent (<5%) level of apoptotic cells in lung tissues from either genotype (our unpublished observations). These data demonstrate that the reduction in air spaces observed in E18.5 KO embryos did not result from increased cell proliferation or compromised cell death.

Lastly, to investigate whether CLASP1 is required for lung maturation, we performed a comparative analysis of histological samples throughout E14.5-PN1 stages of development in WT and KO animals. This revealed a lung developmental delay in *Clasp1* KO mice (Fig. S5) that appears to arise around E15.5-16 and persists until the end of gestation (late pseudo-glandular to early canalicular stages). For instance, the morphology of E16.5-E17.5 KO lungs was equivalent to E15.5-E16.5 WT lungs, with a dense pseudoglandular appearance enclosing a thicker mesenchyme and septa, and with smaller saccules, indicative of poor septal thinning and reduced air-space formation. Taken together, these data support a direct role for CLASP1 in lung maturation.

### In utero glucocorticoid administration only partially overcomes neonatal lethality of Clasp1 KO mice

Glucocorticoids (GCs) are known to play an important role in fetal respiratory development. As so, we wondered whether GC functionality is reduced in the absence of CLASP1. Prenatal GCs, such as dexamethasone, have been used to support fetal lung maturity in pregnancies associated with IUGR and risk of preterm birth (McGoldrick et al., 2020). Therefore, we investigated whether transplacental administration of dexamethasone improves the neonatal outcome in *Clasp1* KO animals. We treated pregnant mice at E16.5 and E17.5 with dexamethasone and allowed the females to deliver. We found that 4/47 KO animals (as opposed to none without dexamethasone) were able to survive for more than 45 min, with 3 KO animals remaining alive for more than 2 h (the animals were then sacrificed for genotyping validation) (Table S1). After the initial (and normal) gasping movements to catch their breath, these KO pups were able to attain a rhythmic respiration, with visible inflation of the lungs and a pulsating area in the thoracic region. Furthermore, around 10-15 min after birth, they were moving and presented a pink color, indicative of blood oxygenation in the organism. Comparative histological examination of dexamethasone-treated lungs from WT and survivor KO animals showed a considerable improvement in the saccular morphology of the *Clasp1* KO lungs, as inferred by the widening of the saccular airspaces (Fig. S6), which were of comparable size as those from WT lungs. However, we noticed that airspace restoration was not uniform in the KO lungs, with few small areas still presenting very small or possibly collapsed saccules. Similar areas were wider and clearly distinguishable in another KO mouse that did not survive after birth, despite dexamethasone treatment (Fig. S6). Overall, these experiments demonstrate that dexamethasone treatment is only partially able to overcome neonatal lethality of *Clasp1* KO mice, suggesting that the observed lung dysfunction is due to additional defects.

### CLASP1 is required for normal lung cell differentiation

To investigate possible mechanisms underlying the lung maturation defects caused by the absence of CLASP1 we analyzed the transcriptomes of three WT and three *Clasp1* KO lungs at E18.5 by RNA-Sequencing (RNA-Seq, Table S2). Principal component analysis (PCA) revealed clustering of the WT and KO samples and indicated that deletion of *Clasp1* does not cause huge genome-wide transcriptional differences (Fig. 6A). Differential gene expression analysis revealed 66 upregulated genes and 112 genes that were significantly downregulated in *Clasp1* KO lungs (Fig. 6B, Table S2).

**Figure 6.**
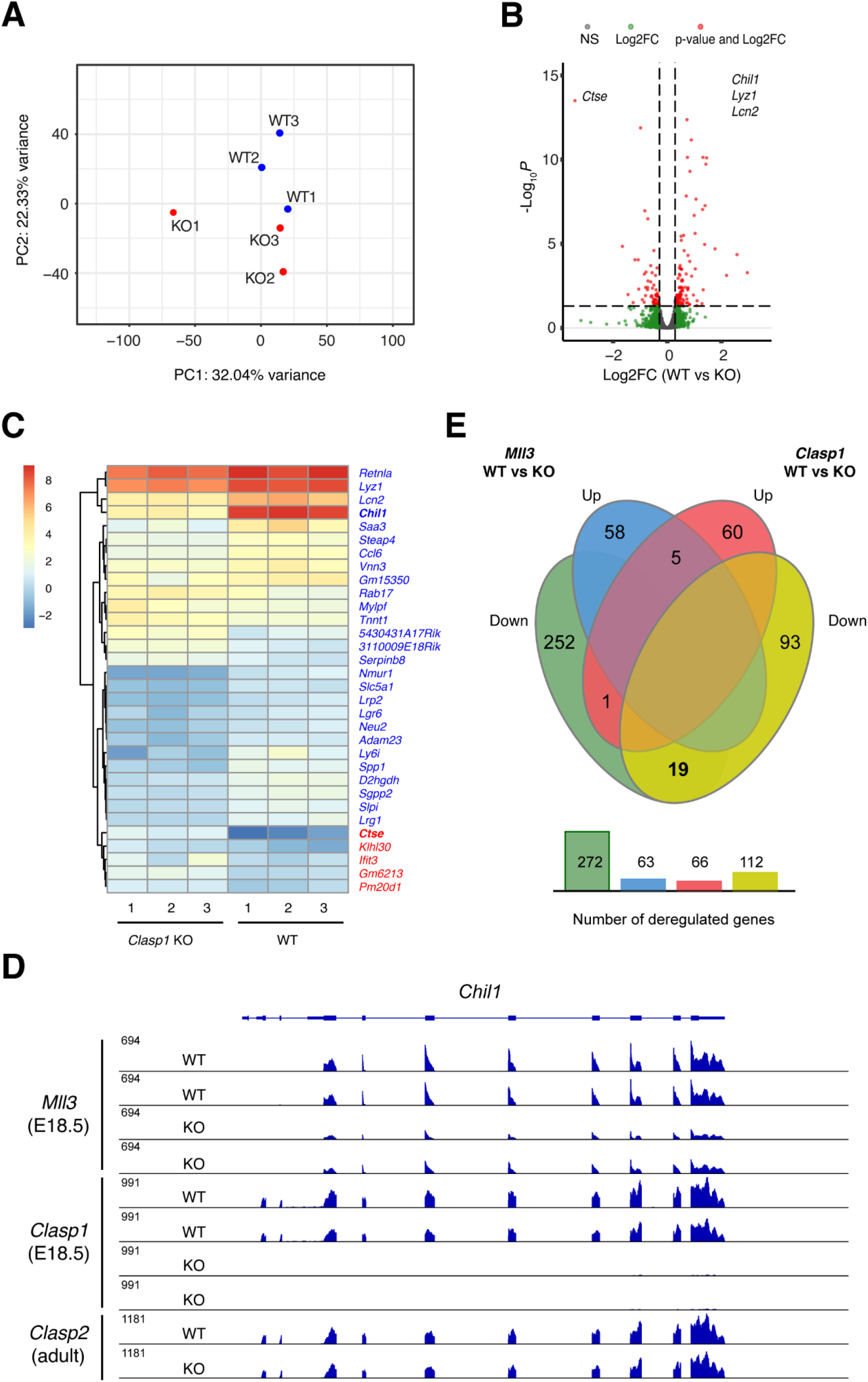
Transcriptome analysis of wild type and *Clasp1* KO lungs. (A) Principal component analysis (PCA). We performed RNA-Seq on mRNA derived from E18.5 lungs from three wild type (WT) and three *Clasp1* KO mice. PCA was performed on the top 10000 most variant genes. (B) Volcano plot of differentially expressed genes. To increase visibility we limited the y-axis value to 15. Hence *Chil1*, *Lyz1*, and *Lcn2* are not visible in the plot. (C) Heatmap showing expression of the most deregulated genes in *Clasp1* KO lungs. These data reveal similar patterns in all three WT samples and all three KO samples. (D) IGV browser image showing the expression of *Chil1* in WT, *Mll3* KO, *Clasp1* KO, and *Clasp2* KO lungs, at E18.5 or in the adult. Both MLL3 and CLASP1 regulate *Chil1* levels, whereas CLASP2 does not. Note that for the *Clasp1* rNA-Seq dataset only two of the three WT and KO RNA-Seq samples are shown. (E) JVENN comparison of DEGs in WT, *Mll3* KO, and *Clasp1* KO lungs at E18.5. Several genes (19 + 5), including *Chil1*, are co-regulated by CLASP1 and MLL3.

Among the most significant differentially expressed genes (DEGs), cathepsin E (*Ctse*) was the highest upregulated one, whereas chitinase-like 1 (*Chil1*), a secreted protein involved in signaling and inflammation (Zhao et al., 2020), was the most down-regulated one (Fig. 6C, Table S2). Indeed, *Chil1* was abundantly expressed in all three WT lungs, yet mRNA levels dropped to almost background in all three *Clasp1* KO lungs (Fig. 6C, D; Table S2). Noteworthy, parallel RNA-Seq analysis of lung samples derived from adult WT and *Clasp2* KO mice suggests that *Chil1* expression is not dependent on CLASP2 (Fig. 6C). Examination of the GEO Omnibus repository revealed another RNA-Seq experiment (GSE146915) carried out in the lungs of E18 WT and *Mll3* KO mice. MLL3 is an H3K4 methyltransferase that is required for lung maturation (Ashokkumar et al., 2020). Interestingly, *Chil1* is downregulated in both *Clasp1* and *Mll3* KO lungs (Fig. 6D), which, overall, showed a significant overlap in DEGs (Fig. 6E). Combined, these data indicate that the same cell types are affected in embryonic lungs of *Mll3* and *Clasp1* KO mice and/or that CLASP1 and MLL3 act in similar pathways to regulate lung maturation.

Importantly, the E18.5 lung RNA-Seq experiment revealed a downregulation of type I cell marker Aquaporin-5 (*Aqp5*) and Aquaporin-1 (*Aqp1*), as well as upregulation of Surfactant protein C (*Sftpc*) (Fig. 7A, Table S2). In contrast to *Sftpc*, Surfactant protein D (*Sftpd*) was downregulated in *Clasp1* KO lungs (Fig. 7A, Table S2), whereas Surfactant protein A1 (*Sftpa1*) and Surfactant protein B (*Sftpb*) were not significantly altered (Table S2). Reduction of *Aqp5* expression was confirmed by quantitative real-time PCR (RT-qPCR) of total RNA isolated from lungs of both genotypes using two different reference genes (Fig. 7B, C). The increased production of Pro-SPC in *Clasp1* KO lungs was also confirmed by immunofluorescence in lung sections and western blot analysis (Fig. 7D-F). Since type II pneumocytes differentiate into type I, and immaturity of type II alveolar cells has been linked to respiratory distress and low rates of neonatal survival (Magnani and Donn, 2020; Mason, 1985), these experiments suggest that facilitated water transport is affected in *Clasp1* KO lungs, and that pneumocyte differentiation is hampered.

**Figure 7.**
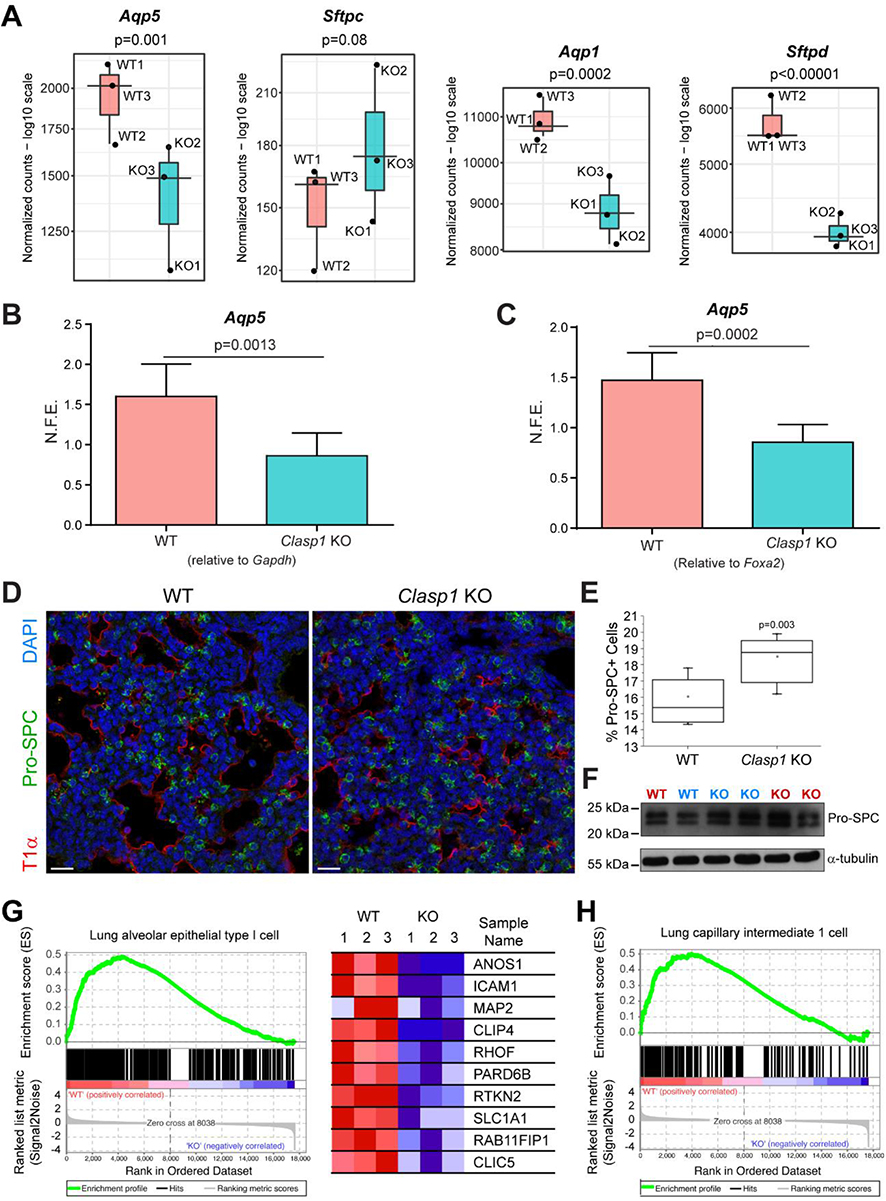
Loss of *Clasp1* alters the ratio of alveolar type I and type II cells in the lungs. (A) RNA-Seq-based expression values of *Aqp1*, *Aqp5*, *Sftpc*, and *Sftpd* in WT and *Clasp1* KO lungs. Values are plotted, with means (lines) and quartiles indicated, and with p values indicated. (B, C) Transcript levels of *Aqp*5 (generalized marker for type I alveolar cells) in *Clasp1* WT and KO lungs. Transcript levels were measured by RT-qPCR and normalized to either *Gapdh* or *Foxa2* levels, expressed as mean ± standard deviation (n=3 for each genotype). (D) Immunofluorescence analysis of E18.5 *Clasp1* WT and KO lungs using type I and type II pneumocyte markers T1a (red) and pro-SPC (green), respectively. DNA was counterstained with DAPI (blue). Scale bar 50 μm. (E) Quantification of pro-SPC-positive cells in *Clasp1* WT and KO lungs (n=3 for each genotype; 9328 and 9369 cells quantified for WT and KO, respectively). (F) Western blot analysis revealing an increase in pro-SPC expression in the KO lungs. Different colours for identify different litters. (G, H) Gene set enrichment analysis (GSEA) of cell type signature gene sets. The RNA-Seq data from three wild type (WT) and three *Clasp1* knockout (KO) mice were used as input for the GSEA. Comparison with the Travaglini dataset (Travaglini et al., 2020) revealed highly significant enrichment of the indicated cell types in WT mice. Expression values of some of the genes in the lung alveolar type I cell data set are listed in (G) to demonstrate differences throughout genotypes.

Lastly, we performed GSEA analysis of cell type signature gene sets, which contain cluster marker genes for cell types identified in single-cell sequencing studies of human tissue. Comparison with a recently established molecular cell atlas of the human lung (Travaglini et al., 2020) revealed enrichment of alveolar epithelial type I cells and lung capillary intermediate cells in WT lungs (Fig. 7G, H), among others, further indicating delayed lung maturation in *Clasp1* KO lungs, particularly of alveolar epithelial type I cells and lung capillary intermediate cells. Overall, we conclude that, in addition to assisting other supporting systems required for normal respiratory function after birth, CLASP1 mediates normal lung cell differentiation and function.

## Discussion

CLASPs are widely conserved proteins expressed in many organisms. Previous work performed in other systems involving the removal or reduction of the single CLASP orthologue resulted in mitotic abnormalities, leading to aneuploidy, polyploidy, infertility and lethality (Inoue et al., 2000; Lemos et al., 2000; Pasqualone and Huffaker, 1994a; Pereira et al., 2006). In mice, just like in humans, there are two paralogues known, CLASP1 and CLASP2 (Akhmanova et al., 2001). Studies on the *in vivo* roles of mammalian CLASPs have thus far all focused on CLASP2; nothing was known about the physiological role of CLASP1. In the present study, we used a classical gene targeting approach to investigate the physiological functions of CLASP1 by creating a *Clasp1*^+/-^ mouse strain able to generate *Clasp1* KO mice that die at birth due to acute respiratory distress.

We were able to demonstrate that, similar to *Clasp2* KO mice (Drabek et al., 2012), *Clasp1* KO mice exhibit diminished fetal growth, including reduced rib cage. Respiratory distress syndrome is a common problem of premature and growth-restricted children. It is also the most important factor concerning neonatal morbidity and mortality, affecting up to 20% of low birth weight children and around 70% of growth-restricted fetuses (Collins et al., 2017). However, no respiratory problems were observed in *Clasp2* KO mice, suggesting that growth restriction *per se* is not the main cause of respiratory distress in *Clasp1* KO mice. Respiratory failure may also occur due to abnormalities in any physiological system that supports breathing. For instance, the respiratory center in the brain controls the rate and depth of respiratory movements of the diaphragm and other respiratory muscles (Guyenet and Bayliss, 2015). *Clasp1* KO mice showed evident microcephaly of unclear significance but our studies on the diaphragm indicated that the innervation pattern detected in the KO muscles was impaired, displaying extended neurites derived from the phrenic nerve. The branching capacity of these neurites was strongly decreased when compared to WT samples. Furthermore, the density of NMJs present at the pre-synaptic side of *Clasp1*-null diaphragms was also significantly reduced. This may impair the mechanical forces generated by the contractile activity of the diaphragm, contributing to the respiratory failure and neonatal lethality observed in *Clasp1* KO mice. Interestingly, CLASP2 has also been implicated in the capture of MTs but at the post-synaptic side of the neuromuscular junction (Schmidt et al., 2012), which in this case was dispensable for neonatal lung function and survival (Drabek et al., 2012).

Necropsy analysis of *Clasp1* KO animals additionally revealed that they suffered from severe atelectasis, raising the possibility of a more direct role of CLASP1 in lung maturation and function. Indeed, we found that, in addition to associated respiratory structures, CLASP1 is expressed in the fetal lung. The maturation of the mammalian lung is a complex process that involves harmonization and synchronization between cell proliferation and differentiation along fetal development, ultimately generating an elaborate organ, composed of several specific cell lineages (Maeda et al., 2007). Our histological observations throughout embryonic and fetal lung development indicate that phenotypic variations between the WT and KO animals arise around E15.5 and are aggravated until birth. The observed reduced saccular expansion associated with an increase in mesenchymal tissue suggests a failure in overall maturation after E15.5, when the transition from early to late canalicular stage is occurring.

Interestingly, the timing of divergence from normal development of the *Clasp1*-null mice overlaps with the known time-course of glucocorticoid production during late murine gestation, which maxes out around E16 (Barlow et al., 1974). Glucocorticoids are key regulators of lung development and are associated with lung maturation, namely terminal differentiation of the distal lung (Bird et al., 2007; Cole et al., 2004; Habermehl et al., 2011). Our data reveal that antenatal glucocorticoid administration could only rescue a small fraction of the KO animals from the striking neonatal mortality observed in the *Clasp1*^-/-^ pups, suggesting other possible causes.

Additional data obtained through RNA-Sequencing, RT-qPCR, western blot and immunofluorescence experiments, further support the hypothesis of generalized impairment of lung maturation after *Clasp1* disruption in mice. Accordingly, E18.5 *Clasp1* KO lungs showed a significant decrease in the transcription of the *Aqp5* and *Aqp1* genes encoding terminal differentiation proteins important for alveolar fluid clearance in neonatal lungs (Song et al., 2000). Moreover, Pro-SPC expression was also increased, further suggesting a pneumocyte maturation delay and hinting for a compromised transport of water between cells in the embryonic lung. Second, using GSEA we compared RNA-Seq datasets of E18.5 WT and *Clasp1* KO lungs with lung cell types identified by single cell RNA-Seq experiments. We observed a highly significant enrichment of lung alveolar epithelial type I cells and lung capillary intermediate 1 cells, and, less significantly, of other cell types, in WT lungs, as inferred from the Travaglini scRNA-Seq dataset (Travaglini et al., 2020). This is consistent with imbalanced lung differentiation of type II into type I cells, as well as the mesenchyme, in the *Clasp1* KO.

A major observation coming out of the RNA-Seq experiments was the complete down-regulation of the *Chil1* gene, encoding Chitinase-like 1, in *Clasp1* KO lungs. This protein (also known as Chitinase 3-like 1 (Chi3l1), YKL-40 or BRP-39) is an important NF-κB-dependent factor and a biomarker for acute and chronic inflammation; it is a secreted glycoprotein and member of the family of chitinase-like proteins (CLPs) that bind to, but do not degrade chitin (He et al., 2013). By screening the GEO Omnibus repository we found that CLASP1 may regulate *Chil1* in conjunction with MLL3, a histone methyltransferase with a specific role in lung development. As hypothesized for the *Mll3* knockout (Ashokkumar et al., 2020), downregulation of common subsets of genes might indicate a reduction in the cell type expressing these genes.

Taken together, our findings support a critical role of CLASP1 in the regulation of cell/tissue growth, bone structure, muscle innervation and pneumocyte differentiation, essential for normal respiratory function and survival at birth. The availability of *Clasp1* KO mice might represent a relevant model system to study developmental disorders associated with respiratory function and growth restriction, and may provide new and better insight into key regulatory pathways underlying lung development in mammals.

## Materials & Methods

### Animals

C57Bl/6 WT mice were obtained from Charles River and Harlan. Animal maintenance and experiments were performed according to protocols that were approved by the Institutional Animal Experimentation Committees at Erasmus University and by the Ethics Committee of Instituto de Biologia Molecular e Celular. The convention for designating mouse ages is such that the morning after conception is embryonic day 0.5 (E0.5) and the day of birth is postnatal day 1 (PN1).

### Targeted inactivation of Clasp1 gene and generation of Clasp1-/- mice

Genomic DNA from 129^ola^ embryonic stem (ES) cells was isolated according to standard procedures (Akagi et al., 1997). The DNA was used as a template for obtaining the two homology regions necessary for the targeting construct. Homology regions were obtained through PCR with Phusion Hot Start High-Fidelity DNA Polymerase (Finnzymes). The 3’end of the first homology region was fused with mCherry and the total fused fragment included, at both ends, a restriction site for ClaI. Two restriction sites for SalI were added to the extremities of the downstream homology region. Fragments were cloned into the ClaI and SalI sites, respectively, of a recipient vector which already contained a loxP-pMC1neor-loxP cassette (Akhmanova et al., 2005). In this construct the neomycin resistance (neo^r^) cassette is transcribed in the antisense direction and is under control of the Polyoma enhancer/TK promoter (PyTK, or PTK). The final targeting construct was verified by sequencing and was subsequently linearized with NotI. The linearized vector was electroporated into E14 129^ola^ ES cells mouse substrain, followed by selection with G418 (200 µg/mL). Individual clones were screened for homologous recombination by Southern blot using a 5’end external probe, as well as by a PCR-based assay for genotyping. Two heterozygous ES cell clones with the correct karyotype were chosen to be injected into C57Bl/6 blastocysts. Chimeric males were mated to C57Bl/6 females to obtain germline transmission of the modified *Clasp1* allele. Heterozygous offspring were maintained on a C57Bl/6 mice background and, simultaneously, crossed between each other in order to generate KO mice. Genotyping was routinely performed by PCR (55°C annealing temperature) using primers:

Gen1: CTCCGCCACTCTCTCCTTTATTGCC

Gen3: CCTCATGTCCTTCCCATAACC

#P261: CGGCATCAGAGCAGCCGATTG

### Southern blot analysis

Genomic DNA was isolated by the sodium dodecyl sulfate-proteinase K procedure (Akagi et al., 1997). DNA was digested with XbaI, blotted onto Hybond H+ membranes (Amersham) and hybridized with a radioactive probe. Hybridization was performed in a rotating hybridizer at 65°C for 24 h in Church Hybmix (0.5 M Na_2_HPO_4_ pH 7.2, 7% SDS, 1 mM EDTA). Membranes were washed extensively at 65°C with Church wash buffer (40 mM Na_2_HPO_4_ pH 7.2, 1% SDS). Hybridization signals were analyzed with a Phosphor Imager (Typhoon Amersham). The following primers were used to generate the radioactive probe for the detection of targeted ES cell clones:

SB_Probe1_F: GCCAGTGAATTTCACTTCCTGG

SB_Probe1_R: GGTTCTCAATCTCCTGTGTACCCAGATGC

### Cell culture and generation of mouse embryonic fibroblasts

Heterozygous mice for *Clasp1* were crossed with each other. E11.5-E13.5 embryos were used to isolate mouse embryonic fibroblasts (MEFs). Blood organs were removed and the head was used to isolate genomic DNA for genotyping. The remaining embryo was finely minced with scissors and incubated for 30 min in Trypsin/EDTA at 37°C. Culture medium, consisting of a mixture 1:1 Dulbecco’s modified eagle medium (DMEM) and Hams F-10 (Invitrogen), supplemented with 10% fetal bovine serum (FBS) and antibiotics, was added and dissociated tissue was transferred to a cell culture dish and cultured at 37°C, 5% CO_2_.

### Western blot

Mice were sacrificed and several organs were collected and frozen in liquid nitrogen. A small amount of each organ was cut and homogenized in ice-cold lysis buffer containing 0.3% Triton X-100 in PBS with protease inhibitor (complete mini EDTA free protease inhibitor, Roche). The suspension was centrifuged for 30 min, 14000 rpm, at 4°C and the protein levels in the supernants were quantified by BSA assays. Whole cell extracts were prepared from E11.5 or E13.5 MEFs according to standard procedures. Briefly, cells were trypsinized when reaching 5−10^6^ confluence, spinned down at room temperature (RT), washed with cold PBS and resuspended in 30-40 µL of lysis buffer containing 0.25 M Tris-HCl pH 7.4, 0.1% Triton X-100 in PBS with protease inhibitor (complete mini EDTA free protease inhibitor, Roche). The samples were incubated on ice for 5 min and centrifuged at 14.000 rpm for 5 min at 4°C and the supernatant was kept for the analysis. All samples were run on 8% sodium dodecyl sulfate-polyacrylamide gel electrophoresis (SDS-PAGE) and electroblotted on nitrocellulose membranes (Protran, Whattman). Blots were blocked in 5% powder milk in TBS pH 7.4 with 0.05% Tween20. Primary antibodies used were rat monoclonal antibodies against CLASP1 (1A6, 1:50-1:100) or CLASP2 (6E3, 1:50-1:100); mouse anti-α-tubulin, clone B512 (1:10.000) (Sigma-Aldrich); PCNA (1:1000) (Cell signaling, #2586); Anti-Prosurfactant Protein C (pro-SP-C) antibody (1:4000) (Millipore, #AB3786). Antibodies were incubated in 2% powder milk in TBS pH 7.4. Signals were detected using the ECL Chemiluminescent Detection System (Amersham). All antibodies were incubated for 1 h at RT. Signal detection was performed with the ECL Chemiluminescent Detection System (Amersham).

### Histology and Immunohistochemistry

Tissues were collected immediately after euthanasia, rinsed 5 –10 min in ice-cold PBS and fixed overnight in 4% fresh formaldehyde in PBS. Three and four µm sections were prepared for HE or immunostaining, respectively. Briefly, all paraffin sections were deparaffinized in xylene and re-hydrated in a graded series of ethanol solutions (100%, 90% and 70%), then rinsed in distilled water. For immunostaining, endogenous peroxidase was blocked with 4% hydrogen peroxide in methanol, for 30-40 min at RT and washed with PBS. Heat-induced antigen retrieval (HIER) was performed by immersing slides into a pre-heated solution containing citrate buffer (10 mM Citric Acid, pH 6.0), microwaved for 5 min at maximum power and cooled down at RT for approximately for 30 min. Unspecific binding was blocked with 3% BSA plus 10% FBS in PBS for 1 h, at RT. Monoclonal Syrian hamster anti-T1α (#8.1 1, 1:10-1:200, Developmental Studies Hybridoma Bank #8.1 1), polyclonal rabbit anti-pro surfactant protein C (1:4000, Millipore, #AB3786), mouse monoclonal acetylated tubulin (Sigma, T7451; 1:20000), mouse anti-smooth muscle actin (Thermo Fisher 1:1200) and anti-α-CC10 (Seven Hills, 1:1000) were used as primary antibodies. Sections were incubated with the primary antibodies overnight, at 4° C. To identify the pro-surfactant protein C and acetylated tubulin, a peroxidase polymer system detection was used after incubation with the primary antibody. T1α, smooth muscle actin and α-CC10 were identified using the respective biotinylated secondary antibodies for 30 min, followed by detection with avidin-biotin complex (ABC, Vector). The reaction was developed using 3-3’-Diaminobenzidine (DAB) (Sigma). For CLASP1 immunostaining, tissues samples were fixed in 4% fresh formaldehyde in PBS and embedded into an OCT (optimal cutting temperature) embedding matrix. Ten μm sections were washed with PBS and blocked for endogenous peroxidase. HIER and blocking solutions were applied as described above. Anti-CLASP1 (1A6, 1:2 or full supernatant) was used, followed by detection with avidin-biotin complex (ABC, Vector).

### Immunofluorescence of lung sections

Formalin-fixed paraffin embedded (FFPE) lung tissue sections with 4 mm were deparaffinised in xylene and rehydrated in a series of ethanol solutions in descending concentrating (100%, 90% and 70% ethanol) until water. Antigen retrieval was performed by boiling tissue sections in citrate buffer (pH 6) for 5 min in a microwave oven, followed by cooling-down at room temperature (RT) for 20 min. Tissue sections were permeabilized with 0.5% Triton X-100 diluted in PBS (0.5% PBS-T) and incubated for 1 h with blocking solution (20% fetal bovine serum diluted in 0.05% PBS-T). Next, tissue sections were incubated overnight (ON) at 4° C in primary antibody solution containing a rabbit antibody against pro-surfactant C (1:4000, Chemicon) and a syrian hamster monoclonal anti-T1α antibody (#8.1 1, 1:10-1:200, Developmental Studies Hybridoma Bank #8.1 1) diluted in blocking solution. After washing (0.05% PBS-T), tissue sections were incubated in the appropriate secondary antibodies (Invitrogen) diluted in blocking solution for 1 h at RT. Nuclei were counterstained with DAPI during incubation with secondary antibody. After washing, tissue auto-fluorescence was quenched with 0.1% Sudan Black diluted in 70% ethanol for 5 min. After thorough washing, coverslips were mounted on microscope slides. Images were acquired by confocal microscopy on a Leica TCS SP5 II.

### Transmission Electron Microscopy

Diaphragms were dissected and fixed by immersion on 2.5% glutaraldehyde (Electron Microscopy sciences, Hatfield, USA) and 2% paraformaldehyde (Merck, Darmstadt, Germany) in cacodylate Buffer 0.1M (pH 7.4), dehydrated and embedded in Epon resin (TAAB, Berks, England). Ultrathin (40–60 nm) sections were prepared on a RMC Ultramicrotome (PowerTome, USA) using diamond knives (DDK, Wilmington, DE, USA) and mounted on 200 mesh copper or nickel grids, stained with uranyl acetate and lead citrate for 5 min each. Afterwards, they were examined under a JEOL JEM 1400 TEM (Tokyo, Japan). Images were digitally recorded using a CCD digital camera Orious 1100W Tokyo, Japan.

### Differential staining of cartilage and bone in whole mouse fetuses by Alcian Blue and Alizarin Red S

Embryos were fixed in 95% ethanol after removal of skin and viscera for 5 days, at RT, in a shaker using orbital agitation. After that time, the ethanol was replaced by acetone and the embryos remained in the shaker with orbital agitation for 2 days. The embryos were rinsed with distilled water. The cartilage was stained by covering the bodies completely with stain solution rocking at RT for 1 day (10 mL per embryo). For every 100 mL, the stain solution contained: 0.4% Alcian blue 8 GX in 70% Ethanol (5 mL), glacial acetic acid (5 mL), 95% ethanol (70 mL), distilled water (20 mL) and 100 µL of 0.5% Alizarin Red (in water) to every 10 mL of stain solution, that was only added at the moment of the staining. The next day, the embryos were briefly rinsed with distilled water and were placed in a graded series of ethanol to remove excess stain: first, they were put overnight in 95% ethanol, followed by 2-3 h in 75% ethanol, 2-3 h in 50% ethanol, 2-3 h in 25% ethanol, and ended with two washes with distilled water. Afterwards, the embryos were transferred to 2% potassium hydroxide for 6 h. After being rinsed in 0.25% potassium hydroxide for 30 min, embryos were cleared in a solution of 20% glycerol and 0.25% potassium hydroxide for 1 h, followed by a solution of 33% glycerol and 0.25% potassium hydroxide for 1 h. Finally, the embryos were cleared in a solution of 50% glycerol and 0.25% potassium hydroxide for 1 h. The embryos were stored in a fresh solution of 50% glycerol and 0.25% potassium hydroxide.

### RNA extraction and transcription analysis by RT-qPCR

Total RNA from mouse embryo lung tissue was obtained by using TRIzol Plus RNA Purification Kit (Ambion, #12183-555) according to the manufacturer’s instructions. Purified total RNA was stored at −80°C, after keeping an aliquot for RNA quality analysys and RNA concentration measurement. For cDNA synthesis, 1 µg of total RNA was transcribed with the iScript Select cDNA Synthesis kit (Bio-Rad) using the random primers and oligo(dTs) supplied, following the manufacturer’s instructions. For real-time PCR of target genes, iQ SYBR Green Supermix (Bio-Rad) was used according to the manufacturer’s instructions. For each analysis, *gapdh* or *foxa2* was used for normalization. RT-qPCRs were performed in the iCycler iQ5 Real-Time PCR Detection System (Bio-Rad Laboratories). The data obtained were analyzed using the Bio-Rad iQ5 Optical System Software v2.1 (Bio-Rad Laboratories).

GAPDH forward primer: CCAGCCTCGTCCCGTAGAC

GAPDH reverse primer: GCCTTGACTGTGCCGTTGA

AQP5 forward primer: CTCCCCAGCCTTATCCATTG

AQP5 reverse primer: TCCTACCCAGAAGACCCATGTG

FoxA2 forward primer: 5’-GGGAGCGGTGAAGATGGA-3’

FoxA2 reverse primer: 5’-TCATGTTGCTCACGGAGGAGTA-3’

### Airway morphometry

Measurements of lung saccular air space were estimated using slides stained with T1α antibody (marker for alveolar type I cells). Microtome sections from stained sections of paraffin embedded mouse lungs (E18.15-E18) were digitally imaged. 3 Random fields of 3 WT and 3 KO embryos were evaluated using an image analysis software, (Image J).

### Motor neuron cultures

Motor neuron cultures were performed as described elsewhere (Conrad et al., 2011). In brief, the thoracic and lumbar portion of the spinal cord was isolated from E12 embryos, and the embryos were genotyped. The individually isolated spinal cords were digested at 37°C for 8 min in 0.25 mg/ml of trypsin (Worthington Biochemical), and the digestion stopped with 10x trypsin-inhibitor (Sigma). After dissociation, the cell suspension was pre-plated into 10 μg/ml lectin (Sigma, L0636) coated 6-well plates for 1 h at RT. After the detachment of the lectin-bounded cells with 30 mM KCl and 0.8% (w/v) NaCl depolarization-solution, cells were counted and plated at 15000 cells/well in 24-well plates with coverslips coated with 0.5mg/ml Poly-DL-ornithine hydrobromide (Sigma, P8638)) and 2.5 μg/ml laminin (Sigma, L2020) or matrigel (BD, 354234), maintained in Neurobasal medium (Invitrogen, 21103) supplemented with 1x B27 (Gibco, 0080085SA), 2 mM L-glutamine (Invitrogen, 25030024), 10 μg/ml CNTF (Peprotech), and 2.5% horse serum (Sigma, H1138).

### Dorsal root ganglia neuron cultures

Dorsal root ganglia were dissected as described elsewhere (da Silva et al., 2014). In brief, dorsal root ganglia were aseptically dissected from E12 embryos, and the embryos genotyped. The isolated ganglia were digested with 0.05% trypsin:EDTA (Invitrogen, 25300) at 37°C for 45 min, and the digestion stopped with HBSS containing 10% FBS. After a 5 min centrifugation at 1000 rpm, the ganglia were resuspended into Neurobasal medium supplemented with 4 g/L of glucose (Sigma), 2 mM L-glutamine, 50 nM NGF (Milipore, 01-125), and 1x B27. After counting the cells, DRG neurons were plated at a cell density of 10000 cells/well in 24-well plates with coverslips coated with matrigel.

### Hippocampal neuron cultures

The hippocampal neuron culture was adapted from a protocol described elsewhere (Leite et al., 2016). Briefly, E17 embryos were individually dissected and genotyped. The individually isolated hippocampi were digested at 37°C for 10 min with 0.06% porcine trypsin solution (Sigma, T4799), the digestion stopped with serum, and the tissue dissociated into single cells. Hippocampal neurons were plated into 24-well plates with poly-L-lysine (Sigma, P0899) coated coverslips at a cell density of 25000 cells/well, and maintained in Neurobasal medium supplemented with 1x B27, 1% penicillin/streptomycin (Invitrogen, 15140-122) and 2 mM L-glutamine.

### Immunofluorescence of cultured neurons

All neurons were fixed with 4% paraformaldehyde in microtubule protection buffer, as described elsewhere (Flynn et al., 2012). Briefly, after 16 and 48 h, cells were fixed for 20 min at RT. After a 5 min permeabilization with 0.2% Triton X-100 at RT, and free aldehydes blocking with 0.2 mM NH4Cl solution for 5 min at RT, the coverslips were blocked for 1 h at RT with 2% BSA (NZYTech, MB04602), 2% FBS (Invitrogen, 10500064), and 0.2% FG (Sigma, G7041) diluted in water. Cells were incubated O/N at 4°C with primary antibodies prepared in 1:10 diluted blocking buffer in PBS, followed by the incubation for 1 h at RT of the respective AlexaFluor-conjugated secondaries. Coverslips were mounted in Fluoroshield with DAPI. Primary antibodies were: acetylated-tubulin (Sigma, T7451; 1:20000); βIII-tubulin (1:5000); all secondary antibodies were used at a dilution of 1:1000.

### Neurite outgrowth evaluation

For neurite outgrowth evaluation, images were taken on an imager Z1 (Carl Zeiss, Germany) with a 20x magnification objective. All images were mounted using Adobe Photoshop CS3 software, and the neurites were evaluated using an automated image analysis routine (SynD) from Matlab. The evaluated parameters were: the length of the longest neurite, the total neurite length, and the number of segments. At least 50 cells were analyzed for each embryo.

### Evaluation of tubulin modifications

For evaluation of tubulin modifications, 65x magnification images were acquired on an Imager Z1 microscope, and the fluorescence intensity at the growth cones was measured for each evaluated tubulin modification and normalized for the βIII-tubulin intensity using the Fiji software. At least 50 cells were analyzed for each embryo.

### Whole-mount tissue immunofluorescence

Muscles at the selected ages were dissected in PBS under a Motic stereomicroscope, fixed with 4% paraformaldehyde (PFA, Sigma) for 6 h at RT, and then storage at 4°C in PBS containing 0.1 M sodium azide (Sigma) until use. The isolated muscles were incubated O/N with 0.1 M glycine (Sigma) at 4°C to block free aldehydes. Tissues were permeabilized for 30 min with 0.5% Triton X-100 (Sigma) at RT, and sequentially incubated at RT for 5 min in 0.2 M NH4Cl (Merck) and 0.1% sodium borohydride (Sigma) to quench auto-fluorescence. After a 1 h blocking of the tissue with 1mg/ml albumin and 0.2% Triton X-100 diluted in PBS at RT, the tissues were incubated for 48 h at 4°C with primary antibodies diluted in the blocking buffer and further incubated with secondary antibodies diluted in blocking buffer for another 48 h. The primary antibody cocktail used was: mouse anti-neurofilament medium (2H3, Developmental Studies Hybridoma Bank; 1:250) and mouse anti-synaptic protein 2 (SV2, Developmental Studies Hybridoma Bank; 1:250). To stain the endplates with fluorescent-labeled BGTx (Molecular Probes), the tissues were further incubated for 1 h at RT with BGTx diluted 1:250 in PBS for whole mounted muscles.

### Quantification of AChR clusters

The number of AChR clusters was quantified using Fiji software in montages of max projections from Z-stacks taken with the fluorescence microscope AxioImager Z1 (Carl Zeiss, Germany) at the central region of the right hemi-diaphragm. The evaluated age was E18, and all the results were normalized for the total area evaluated. Using the same montages, the quantification of the AChR cluster area for the selected area was performed using the crop and magic hand tools from Adobe Photoshop CS3 software.

### Quantification of the innervated muscle area

The evaluation of the axonal extension under the diaphragm was performed using an automated image analysis routine (SynD) from Matlab, in 20x magnification pictures taken with a Axioimager Z1, and mounted using the Adobe Photoshop software.

### Microtubule dynamics

To analyze microtubule dynamics in the growth cone, embryonic hippocampal neurons were isolated from WT and CLASP KO embryos at the embryonic day 16. After 4 days in culture, the hippocampal neurons were transfected with a plasmid encoding for EB3-GFP (Stepanova et al., 2003) using lipofectamine 2000. Twelve hours after transfection, time-lapse recordings were performed as described elsewhere (Pinto-Costa et al., 2020) using a Spinning Disk Confocal System Andor Revolution XD with an iXonEM+DU-897 camera and a IQ 1.10.1 software (Andor Tecnhology). To quantify comet growth speed, the Fiji KymoRescliceWide plugin was used to extract the kymographs, and the start/end positions of the traces defined with the Fiji Cell Counter plugin. In addition, the catastrophe frequency was calculated as described elsewhere (Stepanova et al., 2003).

### RNA-Sequencing and bio-informatics analysis

Total RNA isolation was performed with Trizol-chloroform extraction. After Trizol addition samples were incubated for 5 min at 30°C. Chloroform was added and the aqueous phase was transferred after centrifugation. 100% ethanol was added and RNA was isolated using the RNeasy Mini Kit (Qiagen). RNA sequencing (RNA-Seq) was performed using the Illumina HiSeq platform. RNA-Seq samples were prepared with the Illumina TruSeq Stranded mRNA Library Prep Kit. The resulting DNA libraries were sequenced according to the Illumina TruSeq Rapid v3 protocol on an Illumina HiSeq2000 sequencer, generating single-end reads of 50 bp (43 bp plus 7 bp adapter). Reads were aligned to the mouse genome (mm10 build) using the STAR RNA-Seq aligner (Dobin et al., 2013). Sample scaling and statistical analysis were performed using the R package DESeq2 (Love et al., 2014), within the Octopus-toolkit workflow package (Kim et al., 2018). Principal component analysis (PCA) and gene expression analysis were performed with pcaExplorer (Marini and Binder, 2019). Gene set enrichment analysis was performed using v4.10 of the GSEA software (http://www.gsea-msigdb.org/gsea/index.jsp).

### Statistical analysis

Results were expressed as mean ± standard error of the mean. Comparison data between groups was performed using two-tailed unpaired Student’s t-test or non-parametric Mann-Whitney test. P<0.05 was considered statistically significant.

## Acknowledgements

We thank Cristina Ferrás for input on RT-qPCR, and Rui Fernandes, Francisco Figueiredo and Paula Magalhães for technical help. This research was funded by the Netherlands Organisation for Scientific Research (ZonMW TOP grant 40-00812-98-17045) (to N.G.), as well as Fundação para a Ciência e a Tecnologia of Portugal grants (PTDC/BIA-BCM/112923/2009 and PTDC/MED-ONC/3479/2020), the European Research Council grants PRECISE (260892) and CODECHECK (681443) and La Caixa Health Research Grant (LCF/PR/HR21/52410025) (to H.M.).

## Author contributions

Methodology (ALP, TFS, LTF, MJ, MBK, WFJVL); Investigation, Formal Analysis and Validation (ALP, TFS, LTF, MJ, MBK, WFJVL, NG); Visualization (ALP, TFS, NG, HM); Writing – Original Draft (ALP, HM); Writing – Review and Editing (HM, NG, PB, RR); Supervision (NG, PB, HM); Conceptualization, Project Administration and Funding acquisition (HM, NG).

## References

Akagi, K., V. Sandig, M. Vooijs, M. Van der Valk, M. Giovannini, M. Strauss, and A. Berns. 1997. Cre-mediated somatic site-specific recombination in mice. Nucleic acids research. 25:1766–1773.

Akhmanova, A., C.C. Hoogenraad, K. Drabek, T. Stepanova, B. Dortland, T. Verkerk, W. Vermeulen, B.M. Burgering, C.I. De Zeeuw, F. Grosveld, and N. Galjart. 2001. Clasps are CLIP-115 and -170 associating proteins involved in the regional regulation of microtubule dynamics in motile fibroblasts. Cell. 104:923–935.

Akhmanova, A., A.L. Mausset-Bonnefont, W. van Cappellen, N. Keijzer, C.C. Hoogenraad, T. Stepanova, K. Drabek, J. van der Wees, M. Mommaas, J. Onderwater, H. van der Meulen, M.E. Tanenbaum, R.H. Medema, J. Hoogerbrugge, J. Vreeburg, E.J. Uringa, J.A. Grootegoed, F. Grosveld, and N. Galjart. 2005. The microtubule plus-end-tracking protein CLIP-170 associates with the spermatid manchette and is essential for spermatogenesis. Genes & development. 19:2501–2515.

Ashokkumar, D., Q. Zhang, C. Much, A.S. Bledau, R. Naumann, D. Alexopoulou, A. Dahl, N. Goveas, J. Fu, K. Anastassiadis, A.F. Stewart, and A. Kranz. 2020. MLL4 is required after implantation, whereas MLL3 becomes essential during late gestation. Development. 147.

Barkauskas, C.E., M.J. Cronce, C.R. Rackley, E.J. Bowie, D.R. Keene, B.R. Stripp, S.H. Randell, P.W. Noble, and B.L. Hogan. 2013. Type 2 alveolar cells are stem cells in adult lung. J Clin Invest. 123:3025–3036.

Barlow, S.M., P.J. Morrison, and F.M. Sullivan. 1974. Plasma corticosterone levels during pregnancy in the mouse: the relative contributions of the adrenal glands and foeto-placental units. J Endocrinol. 60:473–483.

Bird, A.D., K.H. Tan, P.F. Olsson, M. Zieba, S.J. Flecknoe, D.R. Liddicoat, R. Mollard, S.B. Hooper, and T.J. Cole. 2007. Identification of glucocorticoid-regulated genes that control cell proliferation during murine respiratory development. J Physiol. 585:187–201.

Boyden, E.A. 1977. Development and growth of airways. In Development of the Lung. W.A. Hodson, editor. Dekker, New York. 3-35.

Cheeseman, I.M., I. MacLeod, J.R. Yates, 3rd, K. Oegema, and A. Desai. 2005. The CENP-F-like proteins HCP-1 and HCP-2 target CLASP to kinetochores to mediate chromosome segregation. Curr Biol. 15:771-777.

Cole, T.J., N.M. Solomon, R. Van Driel, J.A. Monk, D. Bird, S.J. Richardson, R.J. Dilley, and S.B. Hooper. 2004. Altered epithelial cell proportions in the fetal lung of glucocorticoid receptor null mice. Am J Respir Cell Mol Biol. 30:613–619.

Collins, J.J.P., D. Tibboel, I.M. de Kleer, I.K.M. Reiss, and R.J. Rottier. 2017. The Future of Bronchopulmonary Dysplasia: Emerging Pathophysiological Concepts and Potential New Avenues of Treatment. Frontiers in Medicine. 4.

Conrad, R., S. Jablonka, T. Sczepan, M. Sendtner, S. Wiese, and A. Klausmeyer. 2011. Lectin-based isolation and culture of mouse embryonic motoneurons. J Vis Exp.

da Silva, T.F., J. Eira, A.T. Lopes, A.R. Malheiro, V. Sousa, A. Luoma, R.L. Avila, R.J. Wanders, W.W. Just, D.A. Kirschner, M.M. Sousa, and P. Brites. 2014. Peripheral nervous system plasmalogens regulate Schwann cell differentiation and myelination. J Clin Invest. 124:2560–2570.

Desai, T.J., D.G. Brownfield, and M.A. Krasnow. 2014. Alveolar progenitor and stem cells in lung development, renewal and cancer. Nature. 507:190–194.

Dillon, G.M., W.A. Tyler, K.C. Omuro, J. Kambouris, C. Tyminski, S. Henry, T.F. Haydar, U. Beffert, and A. Ho. 2017. CLASP2 Links Reelin to the Cytoskeleton during Neocortical Development. Neuron. 93:1344–1358 e1345.

Dobin, A., C.A. Davis, F. Schlesinger, J. Drenkow, C. Zaleski, S. Jha, P. Batut, M. Chaisson, and T.R. Gingeras. 2013. STAR: ultrafast universal RNA-seq aligner. Bioinformatics. 29:15–21.

Drabek, K., L. Gutierrez, M. Vermeij, T. Clapes, S.R. Patel, J.C. Boisset, J. van Haren, A.L. Pereira, Z. Liu, U. Akinci, T. Nikolic, W. van Ijcken, M. van den Hout, M. Meinders, C. Melo, C. Sambade, D. Drabek, R.W. Hendriks, S. Philipsen, M. Mommaas, F. Grosveld, H. Maiato, J.E. Italiano, Jr., C. Robin, and N. Galjart. 2012. The microtubule plus-end tracking protein CLASP2 is required for hematopoiesis and hematopoietic stem cell maintenance. Cell Rep. 2:781–788.

Drabek, K., M. van Ham, T. Stepanova, K. Draegestein, R. van Horssen, C.L. Sayas, A. Akhmanova, T. Ten Hagen, R. Smits, R. Fodde, F. Grosveld, and N. Galjart. 2006. Role of CLASP2 in microtubule stabilization and the regulation of persistent motility. Curr Biol. 16:2259–2264.

Efimov, A., A. Kharitonov, N. Efimova, J. Loncarek, P.M. Miller, N. Andreyeva, P. Gleeson, N. Galjart, A.R. Maia, I.X. McLeod, J.R. Yates, 3rd, H. Maiato, A. Khodjakov, A. Akhmanova, and I. Kaverina. 2007. Asymmetric CLASP-dependent nucleation of noncentrosomal microtubules at the trans-Golgi network. Developmental cell. 12:917-930.

Fan, G., L. Xiao, L. Cheng, X. Wang, B. Sun, and G. Hu. 2000. Targeted disruption of NDST-1 gene leads to pulmonary hypoplasia and neonatal respiratory distress in mice. FEBS Lett. 467:7–11.

Flynn, K.C., F. Hellal, D. Neukirchen, S. Jacob, S. Tahirovic, S. Dupraz, S. Stern, B.K. Garvalov, C. Gurniak, A.E. Shaw, L. Meyn, R. Wedlich-Soldner, J.R. Bamburg, J.V. Small, W. Witke, and F. Bradke. 2012. ADF/cofilin-mediated actin retrograde flow directs neurite formation in the developing brain. Neuron. 76:1091–1107.

Girao, H., N. Okada, T.A. Rodrigues, A.O. Silva, A.C. Figueiredo, Z. Garcia, T. Moutinho-Santos, I. Hayashi, J.E. Azevedo, S. Macedo-Ribeiro, and H. Maiato. 2020. CLASP2 binding to curved microtubule tips promotes flux and stabilizes kinetochore attachments. J Cell Biol. 219.

Grallert, A., C. Beuter, R.A. Craven, S. Bagley, D. Wilks, U. Fleig, and I.M. Hagan. 2006. S. pombe CLASP needs dynein, not EB1 or CLIP170, to induce microtubule instability and slows polymerization rates at cell tips in a dynein-dependent manner. Genes & development. 20:2421–2436.

Guo, M., Y. Du, J.J. Gokey, S. Ray, S.M. Bell, M. Adam, P. Sudha, A.K. Perl, H. Deshmukh, S.S. Potter, J.A. Whitsett, and Y. Xu. 2019. Single cell RNA analysis identifies cellular heterogeneity and adaptive responses of the lung at birth. Nature Communications. 10:37.

Guyenet, P.G., and D.A. Bayliss. 2015. Neural Control of Breathing and CO2 Homeostasis. Neuron. 87:946–961.

Habermehl, D., J.R. Parkitna, S. Kaden, B. Brugger, F. Wieland, H.J. Grone, and G. Schutz. 2011. Glucocorticoid activity during lung maturation is essential in mesenchymal and less in alveolar epithelial cells. Mol Endocrinol. 25:1280–1288.

He, C.H., C.G. Lee, C.S. Dela Cruz, C.M. Lee, Y. Zhou, F. Ahangari, B. Ma, E.L. Herzog, S.A. Rosenberg, Y. Li, A.M. Nour, C.R. Parikh, I. Schmidt, Y. Modis, L. Cantley, and J.A. Elias. 2013. Chitinase 3-like 1 regulates cellular and tissue responses via IL-13 receptor alpha2. Cell Rep. 4:830–841.

Hur, E.M., Saijilafu, B.D. Lee, S.J. Kim, W.L. Xu, and F.Q. Zhou. 2011. GSK3 controls axon growth via CLASP-mediated regulation of growth cone microtubules. Genes & development. 25:1968–1981.

Inoue, Y.H., M. do Carmo Avides, M. Shiraki, P. Deak, M. Yamaguchi, Y. Nishimoto, A. Matsukage, and D.M. Glover. 2000. Orbit, a novel microtubule-associated protein essential for mitosis in Drosophila melanogaster. The Journal of cell biology. 149:153-166.

Kim, T., H.D. Seo, L. Hennighausen, D. Lee, and K. Kang. 2018. Octopus-toolkit: a workflow to automate mining of public epigenomic and transcriptomic next-generation sequencing data. Nucleic Acids Res. 46:e53.

Kuffer, C., A.Y. Kuznetsova, and Z. Storchová. 2013. Abnormal mitosis triggers p53-dependent cell cycle arrest in human tetraploid cells. Chromosoma. 122:305–318.

Lawrence, E.J., M. Zanic, and L.M. Rice. 2020. CLASPs at a glance. Journal of cell science. 133.

Lee, H., U. Engel, J. Rusch, S. Scherrer, K. Sheard, and D. Van Vactor. 2004. The microtubule plus end tracking protein Orbit/MAST/CLASP acts downstream of the tyrosine kinase Abl in mediating axon guidance. Neuron. 42:913–926.

Leite, S.C., P. Sampaio, V.F. Sousa, J. Nogueira-Rodrigues, R. Pinto-Costa, L.L. Peters, P. Brites, and M.M. Sousa. 2016. The Actin-Binding Protein alpha-Adducin Is Required for Maintaining Axon Diameter. Cell Rep. 15:490–498.

Lemos, C.L., P. Sampaio, H. Maiato, M. Costa, L.V. Omel’yanchuk, V. Liberal, and C.E. Sunkel. 2000. Mast, a conserved microtubule-associated protein required for bipolar mitotic spindle organization. EMBO J. 19:3668–3682.

Logarinho, E., S. Maffini, M. Barisic, A. Marques, A. Toso, P. Meraldi, and H. Maiato. 2012. CLASPs prevent irreversible multipolarity by ensuring spindle-pole resistance to traction forces during chromosome alignment. Nature cell biology. 14:295–303.

Love, M.I., W. Huber, and S. Anders. 2014. Moderated estimation of fold change and dispersion for RNA-seq data with DESeq2. Genome Biol. 15:550.

Maeda, Y., V. Dave, and J.A. Whitsett. 2007. Transcriptional control of lung morphogenesis. Physiological reviews. 87:219–244.

Maffini, S., A.R. Maia, A.L. Manning, Z. Maliga, A.L. Pereira, M. Junqueira, A. Shevchenko, A. Hyman, J.R. Yates, 3rd, N. Galjart, D.A. Compton, and H. Maiato. 2009. Motor-independent targeting of CLASPs to kinetochores by CENP-E promotes microtubule turnover and poleward flux. Curr Biol. 19:1566-1572.

Magnani, J.E., and S.M. Donn. 2020. Persistent Respiratory Distress in the Term Neonate: Genetic Surfactant Deficiency Diseases. Curr Pediatr Rev. 16:17–25.

Maiato, H., E.A. Fairley, C.L. Rieder, J.R. Swedlow, C.E. Sunkel, and W.C. Earnshaw. 2003a. Human CLASP1 is an outer kinetochore component that regulates spindle microtubule dynamics. Cell. 113:891–904.

Maiato, H., A. Khodjakov, and C.L. Rieder. 2005. Drosophila CLASP is required for the incorporation of microtubule subunits into fluxing kinetochore fibres. Nature cell biology. 7:42–47.

Maiato, H., C.L. Rieder, W.C. Earnshaw, and C.E. Sunkel. 2003b. How do kinetochores CLASP dynamic microtubules? Cell Cycle. 2:511–514.

Maiato, H., P. Sampaio, C.L. Lemos, J. Findlay, M. Carmena, W.C. Earnshaw, and C.E. Sunkel. 2002. MAST/Orbit has a role in microtubule-kinetochore attachment and is essential for chromosome alignment and maintenance of spindle bipolarity. J Cell Biol. 157:749–760.

Marini, F., and H. Binder. 2019. pcaExplorer: an R/Bioconductor package for interacting with RNA-seq principal components. BMC Bioinformatics. 20:331.

Mason, R.J. 1985. Pulmonary alveolar type II epithelial cells and adult respiratory distress syndrome. West J Med. 143:611–615.

McGoldrick, E., F. Stewart, R. Parker, and S.R. Dalziel. 2020. Antenatal corticosteroids for accelerating fetal lung maturation for women at risk of preterm birth. Cochrane Database Syst Rev. 12:Cd004454.

Mimori-Kiyosue, Y., I. Grigoriev, G. Lansbergen, H. Sasaki, C. Matsui, F. Severin, N. Galjart, F. Grosveld, I. Vorobjev, S. Tsukita, and A. Akhmanova. 2005. CLASP1 and CLASP2 bind to EB1 and regulate microtubule plus-end dynamics at the cell cortex. The Journal of cell biology. 168:141–153.

Mimori-Kiyosue, Y., I. Grigoriev, H. Sasaki, C. Matsui, A. Akhmanova, S. Tsukita, and I. Vorobjev. 2006. Mammalian CLASPs are required for mitotic spindle organization and kinetochore alignment. Genes Cells. 11:845–857.

Nabhan, A.N., D.G. Brownfield, P.B. Harbury, M.A. Krasnow, and T.J. Desai. 2018. Single-cell Wnt signaling niches maintain stemness of alveolar type 2 cells. Science (New York, N.Y. 359:1118–1123.

Nagase, T., R. Kikuno, K. Ishikawa, M. Hirosawa, and O. Ohara. 2000. Prediction of the coding sequences of unidentified human genes. XVII. The complete sequences of 100 new cDNA clones from brain which code for large proteins in vitro. DNA research : an international journal for rapid publication of reports on genes and genomes. 7:143-150.

Nkadi, P.O., T.A. Merritt, and D.A. Pillers. 2009. An overview of pulmonary surfactant in the neonate: genetics, metabolism, and the role of surfactant in health and disease. Molecular genetics and metabolism. 97:95–101.

Pasqualone, D., and T.C. Huffaker. 1994a. *STU1*, a suppressor of a β-tubulin mutation, encodes a novel and essential component of the yeast mitotic spindle. J Cell Biol. 127:1973–1984.

Pasqualone, D., and T.C. Huffaker. 1994b. STU1, a suppressor of a beta-tubulin mutation, encodes a novel and essential component of the yeast mitotic spindle. The Journal of cell biology. 127:1973–1984.

Pereira, A.L., A.J. Pereira, A.R. Maia, K. Drabek, C.L. Sayas, P.J. Hergert, M. Lince-Faria, I. Matos, C. Duque, T. Stepanova, C.L. Rieder, W.C. Earnshaw, N. Galjart, and H. Maiato. 2006. Mammalian CLASP1 and CLASP2 cooperate to ensure mitotic fidelity by regulating spindle and kinetochore function. Mol Biol Cell. 17:4526–4542.

Pinto-Costa, R., S.C. Sousa, S.C. Leite, J. Nogueira-Rodrigues, T. Ferreira da Silva, D. Machado, J. Marques, A.C. Costa, M.A. Liz, F. Bartolini, P. Brites, M. Costell, R. Fassler, and M.M. Sousa. 2020. Profilin 1 delivery tunes cytoskeletal dynamics toward CNS axon regeneration. J Clin Invest. 130:2024–2040.

Ramirez, M.I., G. Millien, A. Hinds, Y. Cao, D.C. Seldin, and M.C. Williams. 2003. T1alpha, a lung type I cell differentiation gene, is required for normal lung cell proliferation and alveolus formation at birth. Dev Biol. 256:61–72.

Sayas, C.L., S. Basu, M. van der Reijden, E. Bustos-Moran, M. Liz, M. Sousa, I.W.F.J. van, J. Avila, and N. Galjart. 2019. Distinct Functions for Mammalian CLASP1 and -2 During Neurite and Axon Elongation. Frontiers in cellular neuroscience. 13:5.

Schmidt, N., S. Basu, S. Sladecek, S. Gatti, J. van Haren, S. Treves, J. Pielage, N. Galjart, and H.R. Brenner. 2012. Agrin regulates CLASP2-mediated capture of microtubules at the neuromuscular junction synaptic membrane. The Journal of cell biology. 198:421–437.

Shahbazi, M.N., D. Peña-Jimenez, F. Antonucci, M. Drosten, and M. Perez-Moreno. 2017. Clasp2 ensures mitotic fidelity and prevents differentiation of epidermal keratinocytes. Journal of cell science. 130:683–688.

Song, Y., N. Fukuda, C. Bai, T. Ma, M.A. Matthay, and A.S. Verkman. 2000. Role of aquaporins in alveolar fluid clearance in neonatal and adult lung, and in oedema formation following acute lung injury: studies in transgenic aquaporin null mice. J Physiol. 525 Pt 3:771–779.

Stepanova, T., J. Slemmer, C.C. Hoogenraad, G. Lansbergen, B. Dortland, C.I. De Zeeuw, F. Grosveld, G. van Cappellen, A. Akhmanova, and N. Galjart. 2003. Visualization of microtubule growth in cultured neurons via the use of EB3-GFP (end-binding protein 3-green fluorescent protein). J Neurosci. 23:2655–2664.

Travaglini, K.J., A.N. Nabhan, L. Penland, R. Sinha, A. Gillich, R.V. Sit, S. Chang, S.D. Conley, Y. Mori, J. Seita, G.J. Berry, J.B. Shrager, R.J. Metzger, C.S. Kuo, N. Neff, I.L. Weissman, S.R. Quake, and M.A. Krasnow. 2020. A molecular cell atlas of the human lung from single-cell RNA sequencing. Nature. 587:619–625.

Turgeon, B., and S. Meloche. 2009. Interpreting neonatal lethal phenotypes in mouse mutants: insights into gene function and human diseases. Physiological reviews. 89:1–26.

Warburton, D., J. Gauldie, S. Bellusci, and W. Shi. 2006. Lung development and susceptibility to chronic obstructive pulmonary disease. Proceedings of the American Thoracic Society. 3:668–672.

Whitsett, J.A., T.V. Kalin, Y. Xu, and V.V. Kalinichenko. 2019. Building and Regenerating the Lung Cell by Cell. Physiological reviews. 99:513–554.

Whitsett, J.A., S.E. Wert, and T.E. Weaver. 2010. Alveolar surfactant homeostasis and the pathogenesis of pulmonary disease. Annual review of medicine. 61:105–119.

Wittmann, T., and C.M. Waterman-Storer. 2005. Spatial regulation of CLASP affinity for microtubules by Rac1 and GSK3beta in migrating epithelial cells. The Journal of cell biology. 169:929–939.

Zhao, T., Z. Su, Y. Li, X. Zhang, and Q. You. 2020. Chitinase-3 like-protein-1 function and its role in diseases. Signal Transduct Target Ther. 5:201.

