## Supplementary Figures and Table S1 for "CLASP1 is essential for neonatal lung function and survival in mice"

### SUPPLEMENTARY FIGURES AND TABLES

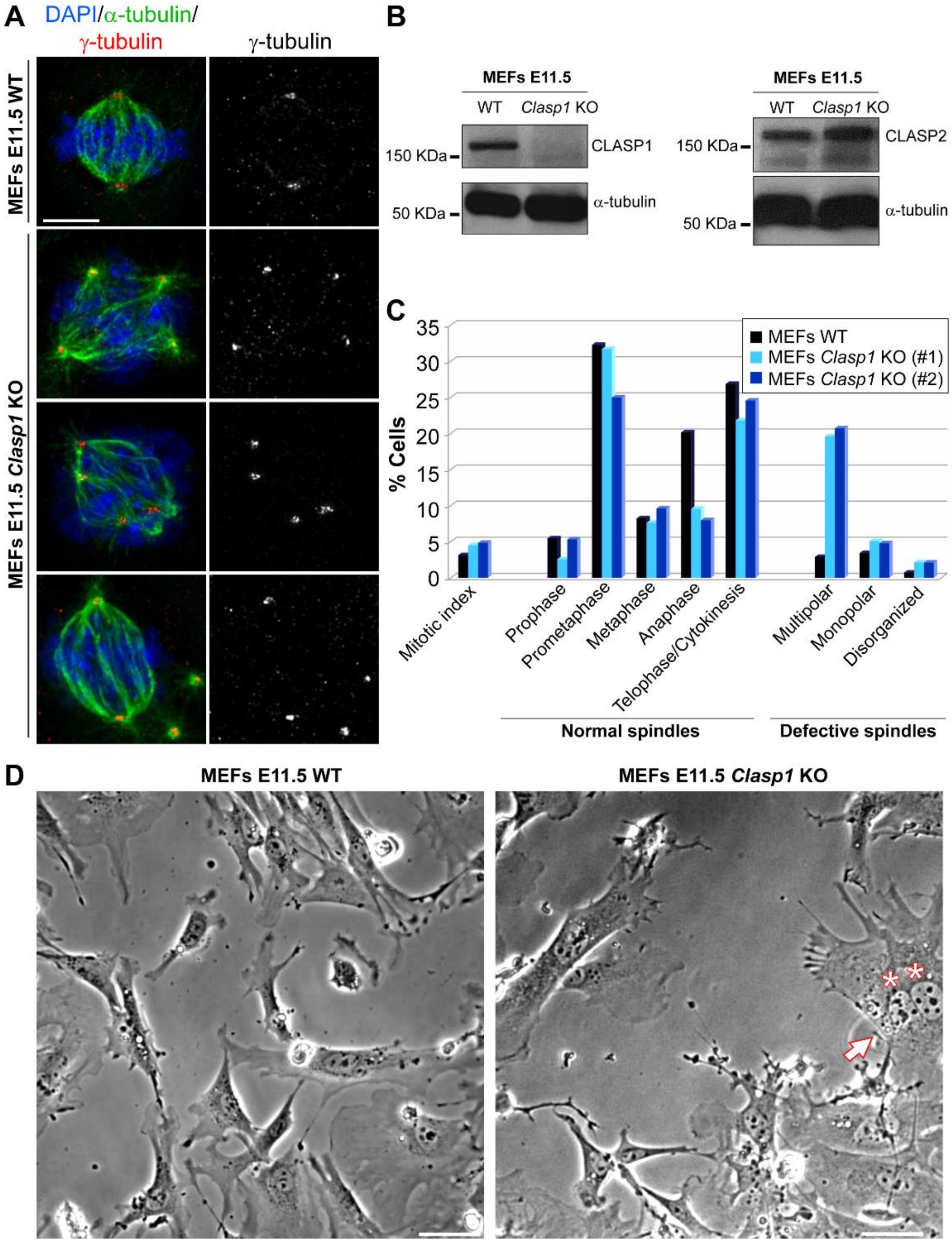


**Figure S1. Incidental cell division defects in mouse embryonic fibroblasts derived from *Clasp1* KO mice, Related to Figure 1.** (A) Multipolar spindle defects in *Clasp1* KO MEFs observed by immunofluorescence analysis with α-tubulin (microtubules; green), γ-tubulin (centrosomes; red) antibodies. DNA was counterstained with DAPI (blue). Scale bar is 5 μm. (B) Western blot analysis confirming the absence of CLASP1 protein (but not CLASP2) from the *Clasp1* KO MEFs (E11.5). (C) Quantification of mitotic phenotypes in two independent MEF lines from *Clasp1* KO animals. Note the slight increase in mitotic index and clear reduction of anaphase cells in KO MEFs, which showed also a high frequency of multipolar spindles. (D) Snapshot from live-cell phase-contrast microscopy analysis of WT and *Clasp1* KO MEFs (E11.5) showing an example of a binucleated cell (red arrow). The two nuclei are indicated (red asterisks). Scale bar 50 μm.

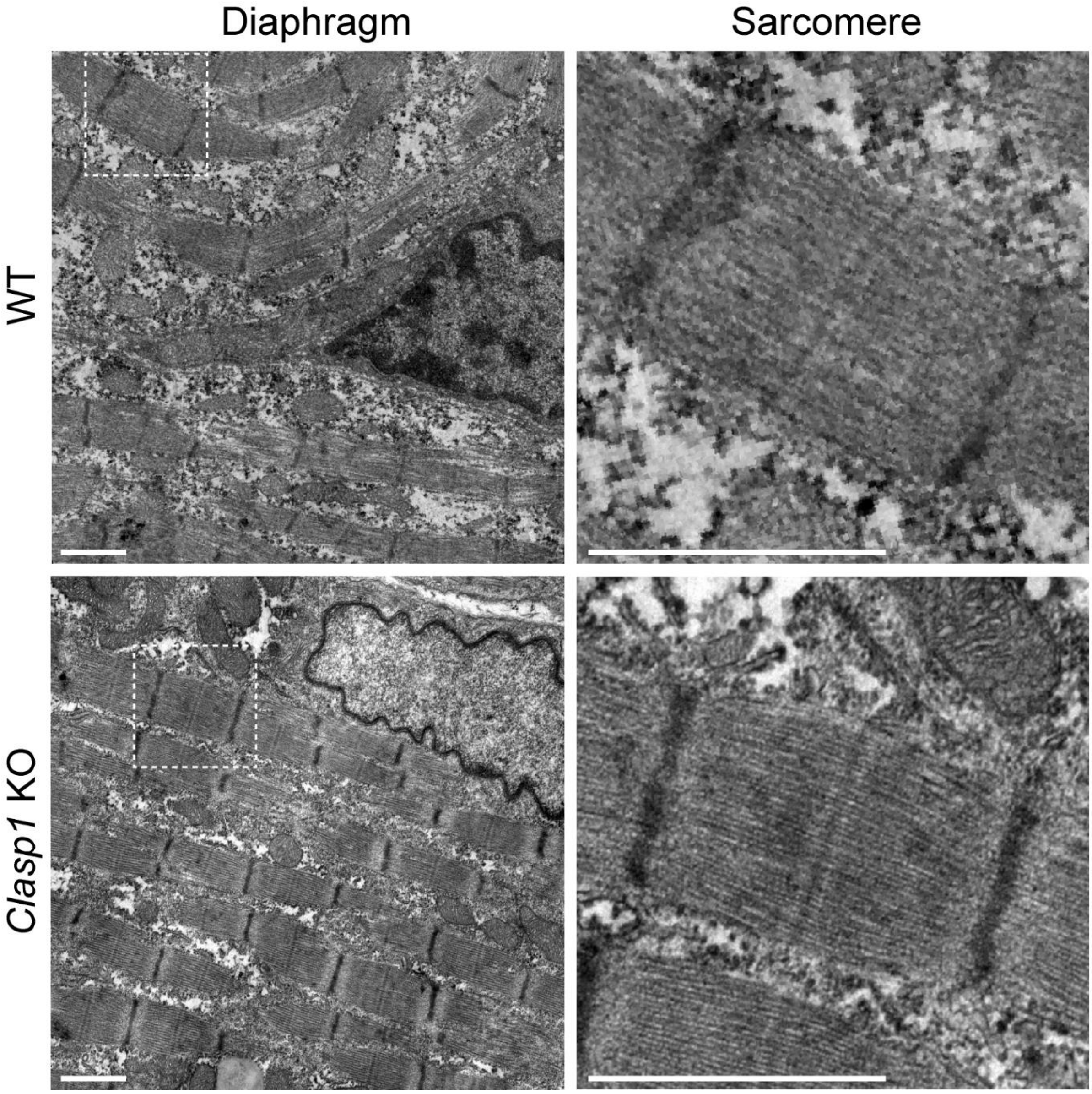


**Figure S2. Ultra-structural analysis of Diaphragms, Related to Figure 3.** Representative ultrathin section of diaphragms from WT and *Clasp1* KO mice showing normal sarcomere architecture (zoom, right panels). Scale bar is 1 μm.

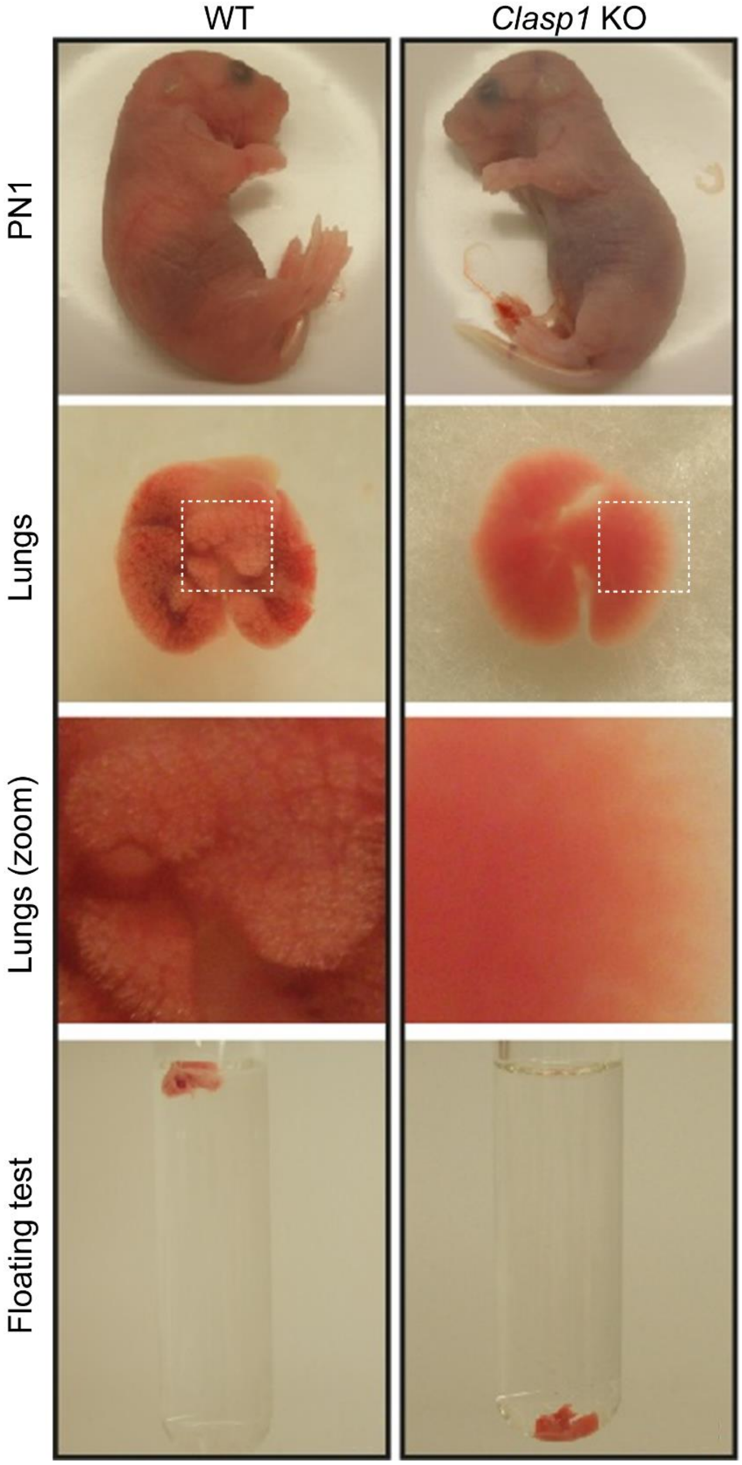


**Figure S3. Newborn *Clasp1* KO lungs show a drastic reduction in air inflation, Related to Figure 5.** Wild type (WT) and *Clasp1* KO newborn mice with the respective lung macroscopic features and water-floating test result. Note the typical cyanotic colour of KO pups relative to the pinkish colour of WT littermates (top). The foamy aspect of the WT lungs is missing in KO lungs, indicating the absence of air inside the alveoli. At the bottom, the water-floating test confirm the lung collapse of KO animals, whose lung samples immediately sunk upon release in the saline solution.

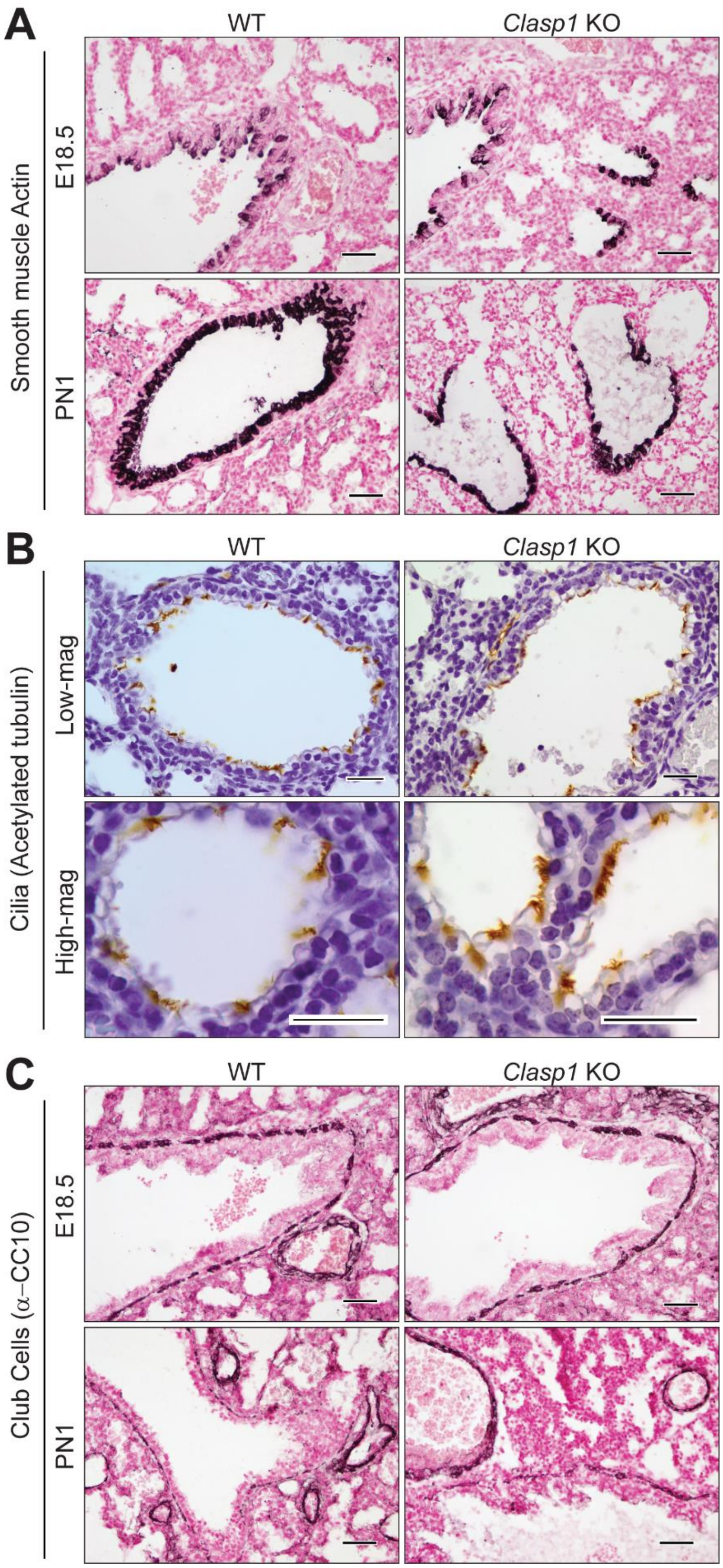


**Figure S4. Histo-morphological analysis of the developing lung throughout embryonic development (E14.5-PN1), Related to Figure 5.** HE staining of lung sections from WT and *Clasp1* KO animals collected from timed pregnancies, with indicated developmental stages. Scale bars 150 µm.

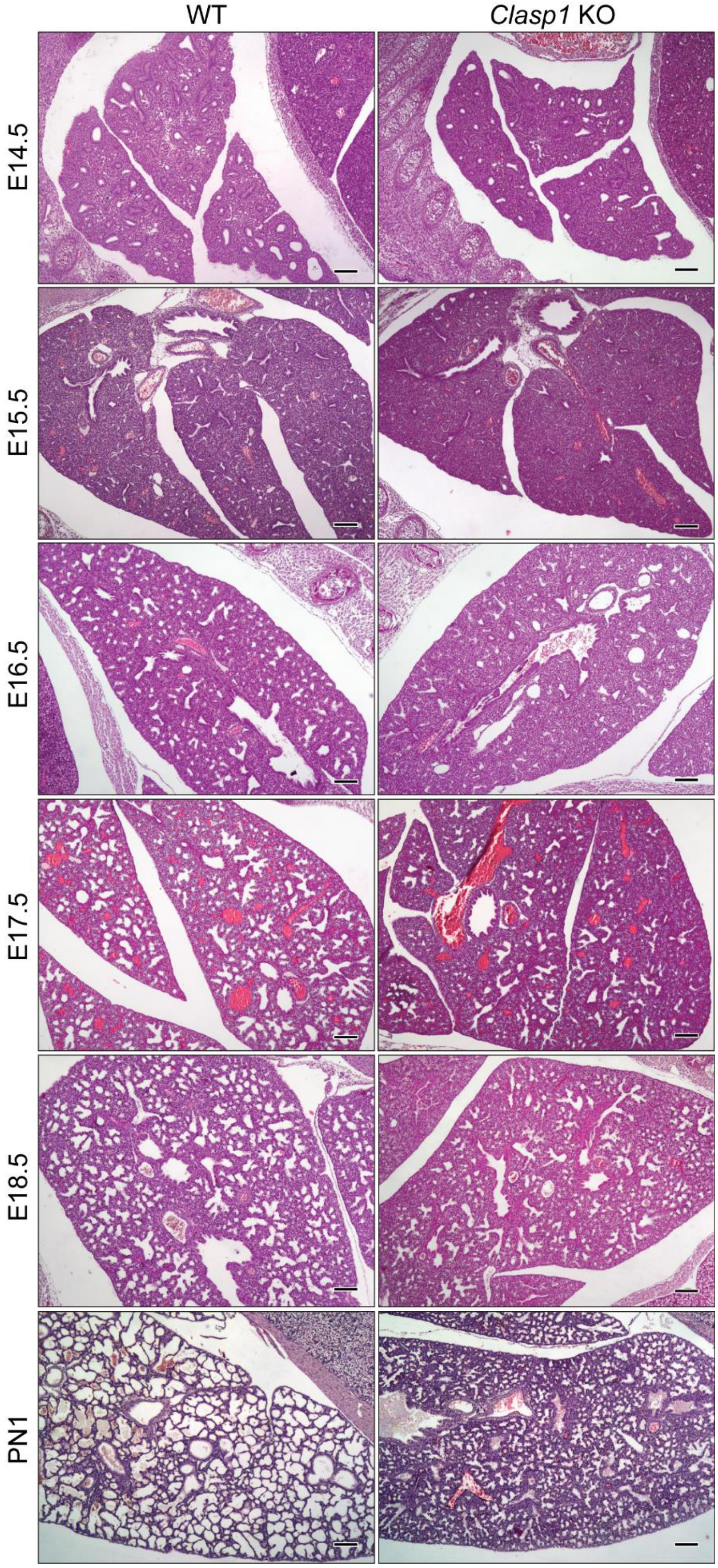


**Figure S5. Cellular analysis of late stage gestation mouse lungs, Related to Figure 5.** Histological sections of WT and *Clasp1*-null lungs stained with (A) α-SMA, (B) α-acetylated tubulin, and (C) α-CC10 antibodies. Scale bar 25 µm.

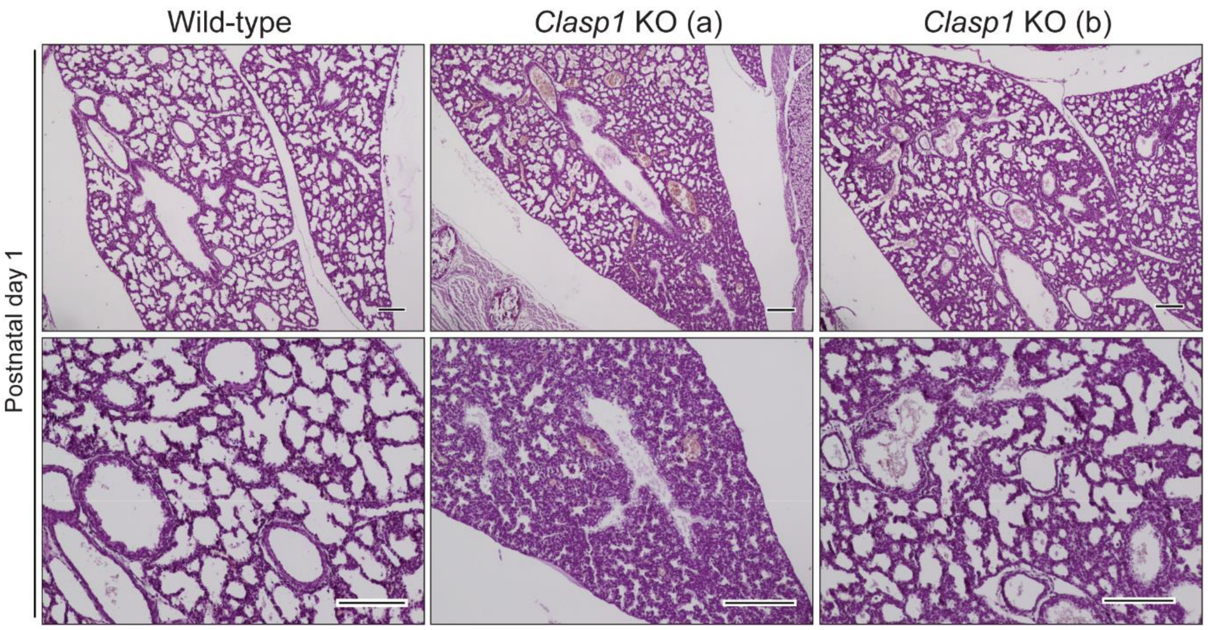


**Figure S6. Exogenous administration of glucocorticoids promotes lung maturation and partially rescues postnatal lethality, Related to Figure 5.** (A) Representative light microscopic images from wild-type and *Clasp1* KO lung paraffin sections stained with HE, after being submitted to a dexamethasone treatment, at different magnifications.

**Table S1.** Survival results from *in utero* dexamethasone administration.

| | Alive | Dead | Total |
|---|---|---|---|
| WT | 51 | 1 | 52 |
| HT | 63 | 3 | 66 |
| *Clasp1* KO | 3 | 43 (+1) | 47 |

**Table S2.** RNA-Sequencing and differential gene expression analysis from E18.5 lungs derived from wild type and *Clasp1* KO embryos (see accompanying excel file)
